# It takes two: A Widespread Temperate Bacteriophage Contributes to Regulation of the Type III Secretion System in *Pseudomonas syringae*

**DOI:** 10.64898/2026.07.06.736757

**Authors:** Daniel Maddock, Sophia Liberto, Brooke Ognian, George W. Sundin, Michelle T. Hulin

## Abstract

- The *Pseudomonas syringae* species complex includes major crop pathogens that use a type III secretion system (T3SS) to inject effectors into plant cells, suppressing immunity and promoting disease.
- The cherry canker pathogen *Pseudomonas amygdali* pv. *morsprunorum* (Pam) carries the effector gene *hopAR1* on a prophage, PamPP1, which belongs to a novel Caudoviricetes family widespread across the *P. syringae* complex and likely acquired before pathovar divergence.
- Deletion of PamPP1 shows that this prophage enhances Pam virulence independently of hopAR1, instead it alters the T3SS operon expression both *in vitro* and *in planta*.
- These prophage-driven transcriptional changes likely reshape Pam’s interaction with plant immunity, highlighting how bacteriophages rewire bacterial transcriptomes and contribute to the evolution and emergence of plant diseases.

## Introduction

Bacteria are pathogens of a wide range of eukaryotic organisms. As such, much effort has been spent researching virulence mechanisms and the evolution of these. Mobile genetic elements (MGEs), including plasmids, integrative conjugative elements, transposons, and bacteriophages (phages) are key features of bacterial evolution (Maddock & Hulin, 2025). Phages are viruses of bacteria and are ubiquitous across diverse environments. Lytic phages infect and immediately produce new phage particles and lyse their host. By comparison, temperate phages infect and enter the lysogenic cycle, where they integrate their genome into the bacterial host’s genome and become dormant (termed a prophage) (Zhang *et al*., 2022). Temperate phages can switch from the lysogenic cycle to the lytic cycle to leave and infect new hosts. During this process, they can acquire bacterial genes. Genes involved in fitness in particular environments, such as virulence or antimicrobial resistance genes, are often acquired by prophages and transferred to new hosts driving bacterial evolution (Knowles *et al*., 2016; Inglis *et al*., 2025). Prophages are incredibly common, with one study showing 75% of 13,000 bacterial genomes contained prophages (López-Leal *et al*., 2022).

Prophages can alter bacterial host gene expression. In the citrus pathogen *Candidatus* Liberibacter asiaticus, transcriptional regulation was among processes affected by prophage integration (Yin *et al*., 2025). In *Escherichia coli,* a prophage encodes a transcription factor YdaT which alters expression of the regulator RcsA to impact adhesion, motility and exopolysaccharide synthesis (Wons *et al*., 2025). *Clostridium difficile* contains a prophage gene encoding RepR which downregulates the production of toxins (Govind *et al*., 2009). These phenotypic changes, together with the role of phages in controlling population dynamics (causing 20-40% of bacterial lysis) indicate phages heavily influence bacterial evolution (Chevallereau *et al*., 2022).

The *Pseudomonas syringae* species complex (PSSC) encompasses diverse plant pathogenic and environmental strains with global distribution. The complex is divided into 13 phylogenetic groups (phylogroups) and over 60 pathogenic variants (pathovars) classified based on the plant host they infect (Baltrus *et al*., 2016). PSSC strains collectively cause disease on over 500 plant species, with major relevance to agriculture (López-Pagán *et al*., 2025). They frequently live epiphytically on the plant surface prior to entering the apoplast to cause disease (Macho *et al*., 2007). Most *P. syringae* strains possess a Hrp1 T3SS which they utilize to inject T3Es into plant cells (“Dynamic Evolution of Pathogenicity Revealed by Sequencing and Comparative Genomics of 19 Pseudomonas syringae Isolates | PLOS Pathogens”). When *P. syringae* infects the plant host, conserved molecules such as flagellin, termed pathogen-associated molecular patterns (PAMPs), are recognized by pattern-recognition receptors to initiate PAMP-triggered immunity (PTI). To suppress PTI, T3Es act to inhibit immune processes. However, plants can perceive T3Es directly or indirectly using R proteins, which initiate effector-triggered-immunity (ETI) (Block & Alfano, 2011). If PTI/ETI is triggered, expression of pathogenesis-related (PR) proteins is activated by salicylic acid accumulation and Systemic Acquired Resistance (SAR) pathways are triggered for long-distance induced resistance (Vlot *et al*., 2021; dos Santos & Franco, 2023).

Within phylogroup 3 of the PSSC, the species *P. amygdali* encompasses a wide range of hemi-biotrophic host-specialized pathogens of woody hosts (Berge *et al*., 2014). Pathovar *morsprunorum* (Pam) (also known as *P. syringae* pv. *morsprunorum* race 1) causes bacterial canker disease of stone fruit (*Prunus*) species such as sweet cherry (Gilbert *et al*., 2010). Pam lives epiphytically within the phyllosphere before moving through natural openings and wounds to colonize the apoplast (Hulin *et al*., 2018b). Symptoms of the disease on leaves appear as necrotic leaf spots, water-soaked lesions on fruit, and cankers involving necrosis and gummosis of woody tissues (Hulin *et al*., 2020). Previous analysis showed that the T3E gene *hopAR1* is present in Pam within a prophage region (Hulin *et al*., 2018a). We demonstrated this prophage (PamPP1) can excise, circularize, and mobilize into a distantly related *P. syringae* phylogroup 10 strain when co-inoculated with Pam on the leaf surface (Hulin *et al*., 2023b).

The movement of phages within epiphytic bacterial communities indicates that the plant surface is a conducive environment for Horizontal Gene Transfer (HGT). The presence of the virulence-related genes on these phages provides a model for observing the ecological impact of HGT and its contribution to disease emergence. Here we set out to examine several different questions. What is the distribution of the PamPP1 prophage in the *P. amygdali* clade and the wider PSSC? Finally, does the PamPP1 prophage alter virulence of Pam and thus play a role in disease emergence?

## Methods

### Bacteria culturing, plasmids and primers

Strains are listed in Table 1, plasmids and primers are in Tables S1 and S2. Kings Medium B (KB) and Lysogeny Broth (LB) were used for standard culturing (King *et al*., 1954; “Sambrook, J., Fritsch, E. R., & Maniatis, T. (1989). Molecular Cloning A Laboratory Manual (2nd ed.). Cold Spring Harbor, NY Cold Spring Harbor Laboratory Press. - References - Scientific Research Publishing”). To mimic conditions within the plant apoplast, hypersensitive response and pathogenicity (hrp)-inducing minimal medium (HIM) was used (Huynh *et al*., 1989). The following antibiotics were used: kanamycin (50 µg/mL), rifampicin (50 µg/mL), nitrofurantoin (200 µg/mL), cycloheximide (25 µg/mL).

**Table 1.**
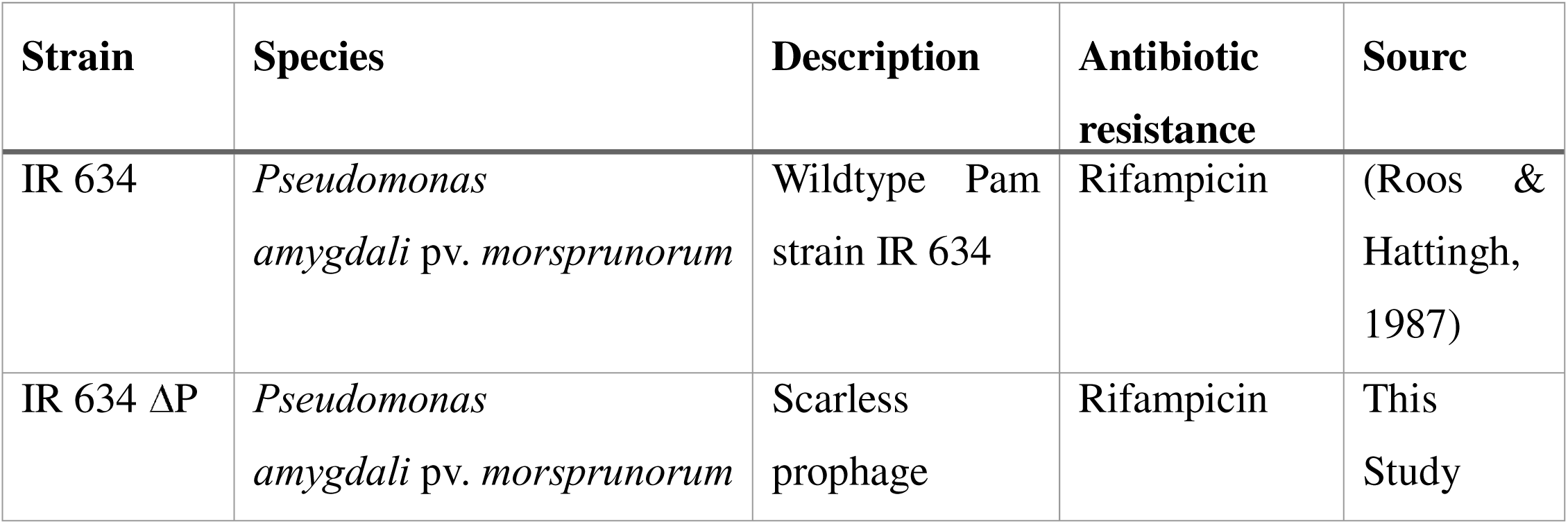
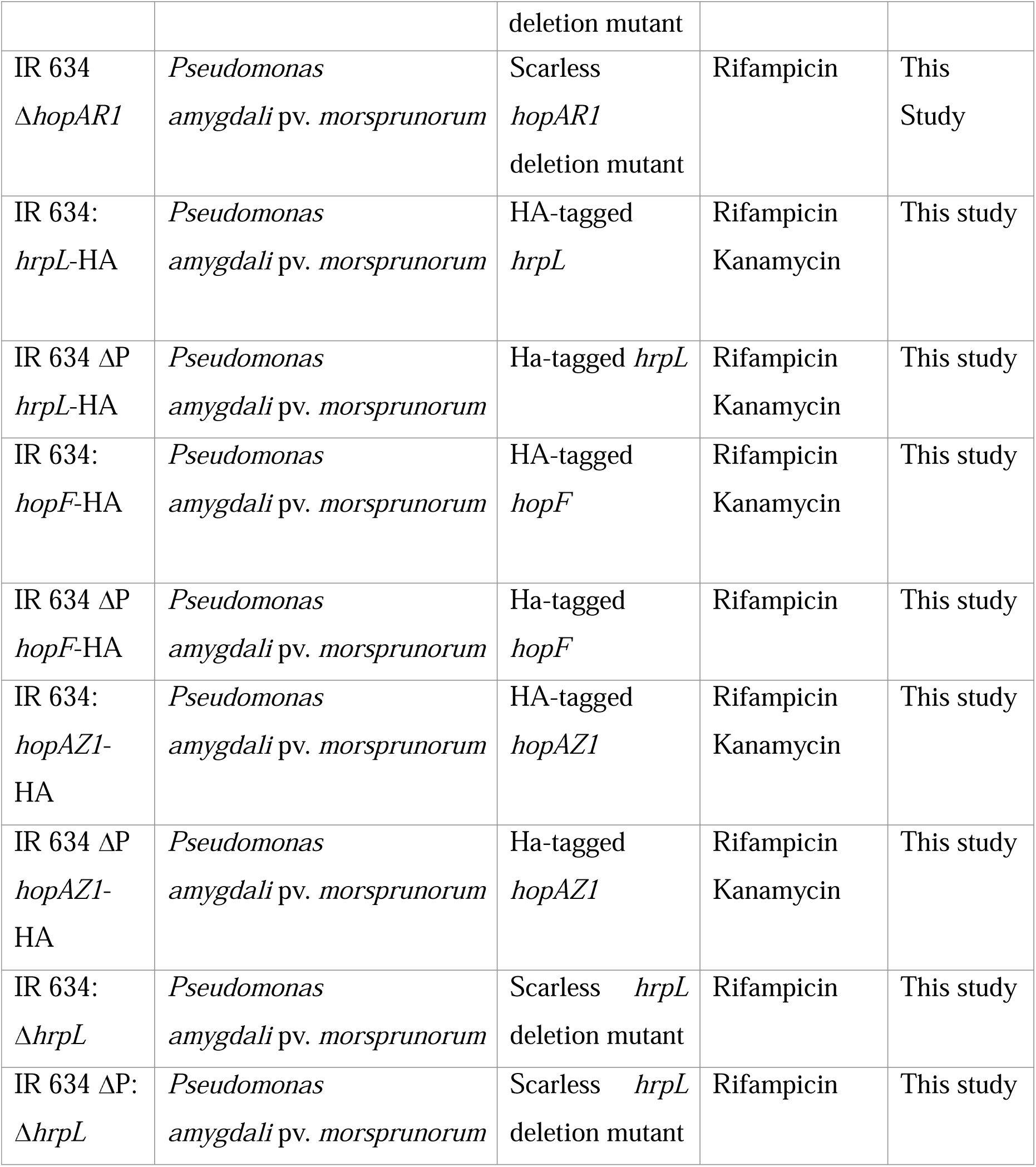
List of *P. syringae* strains used in this study.

### RNA sequencing

RNA sequencing was performed for Pam grown *in vitro.* Overnight KB cultures were centrifuged at 17,000 RPM for 5 mins and washed twice in 10mM MgCl_2_ to remove antibiotics, and adjusted to an OD_600_ of 0.2. Samples were centrifuged at max speed and resuspended in HIM and grown for 25 h. 2 mL aliquots were taken at specific time points (0, 5 and 25hr) and cells pelleted by centrifugation at max speed for 5 mins. Pellets were frozen in liquid nitrogen and stored at –80°C. RNA extractions were performed using Trizol and chloroform, followed by Qiagen RNeasy kit with on-column DNase digestion with an additional removal with the Turbo DNA-Free kit (Invitrogen, USA). RNA was sent for paired-end Illumina sequencing with Novogene (NovaSeq X Plus Series (PE150) (2 Gb raw data per sample)). Raw data are available in the NCBI sequence read archive accession PRJNA1419900.

### Bioinformatics

A total of 2542 PSSC genomes (taxid136849) were downloaded in April 2024. These were filtered to 1669 genomes, removing those of lower quality and reducing the number of clonal genomes as done previously (Hulin *et al*., 2023a). Genomes were annotated with Bakta v.1.9.4 (Schwengers *et al*., 2021). Panaroo v1.5.2 (Tonkin-Hill *et al*., 2020) was utilized to generate a core genome alignment. Veryfastree v4.0.5 (“Very Fast Tree: speeding up the estimation of phylogenies for large alignments through parallelization and vectorization strategies | Bioinformatics | Oxford Academic”) generated a maximum-likelihood phylogenetic tree using the generalized-time-reversible model with 100 bootstrap replicates. Prophage sequences were predicted using PhiSpy v4.2.1 (Akhter *et al*., 2012) and PHASTEST (Wishart *et al*., 2023). Phage taxonomic classification used vConTACT3 v3.1.6, database v230 (Bolduc *et al*., 2025). Phages were clustered with 1283 other *Pseudomonas* phages obtained from the Virus-Host database (https://www.genome.jp/ftp/db/virushostdb/) using VipTreeGen v1.1.3 (Nishimura *et al*., 2017) to produce a neighbor-joining tree. This was input into R v4.3.2 (“R Core Team (2021) R A Language and Environment for Statistical Computing. R Foundation for Statistical Computing, Vienna. - References - Scientific Research Publishing”) to perform hierarchical clustering to generate groups based on genetic similarity.

For analysis of the *Pseudomonas amygdali* clade, Panaroo core genome alignments were input to IQ-TREE2 (Minh *et al*., 2020). Prophages were annotated with Pharokka (v1.7.3) (Bouras *et al*., 2023) using a custom T3 effector (T3E) HMM database of the *Pseudomonas syringae* Type III Effector Compendium (Laflamme *et al*., 2020). Custom databases were made from multiple sequence alignments of each effector family (clustal-omega v1.2.4) (Sievers *et al*., 2011) using HMMBUILD (HMMER v3.4) (Eddy, 08/2023). Further gene function prediction was performed using HHpred https://toolkit.tuebingen.mpg.de/tools/hhpred (Söding *et al*., 2005). For PamPP1 prophages, genomic regions with an additional 20 Kb on either side were extracted and annotated, and PHASTEST used to confirm prophage completeness. Regions were aligned using clinker (v0.0.29) (“clinker & clustermap.js: automatic generation of gene cluster comparison figures | Bioinformatics | Oxford Academic”).

For RNA-seq analysis, data were assessed with FastQC (v0.12.0) (“Andrews, S. (2010) FastQC A Quality Control Tool for High Throughput Sequence Data. - References - Scientific Research Publishing”) and underwent adapter trimming, filtering, and pre-processing using fastp (v1.0) (“fastp 1.0: An ultra fast all round tool for FASTQ data quality control and preprocessing - Chen - 2025 - iMeta - Wiley Online Library”). QC outputs were assessed and rRNA removed using ribodetector (v0.3.1) (Deng *et al*., 2022). Cleaned reads underwent transcript alignment and quantification against a generated transcriptome file concatenated with a decoy file using Salmon (v1.10.1) (Patro *et al*., 2017). Transcript quantifications were analyzed using DESeq2 (Love *et al*., 2014). Gene networks were visualized with Cytoscape (v3.10.4) (Shannon *et al*., 2003) using string enrichment against a custom database made using the proteome of Pam IR 634 (organism ID: STRG0A01MPX made from the Pam IR 634 genome (JBTVRL000000000.1) (Szklarczyk *et al*., 2025). Heatmaps and upset plots of normalized read counts were generated using TBtools-II (Chen *et al*., 2023). Figures and statistical analysis were generated using Rstudio (v4.4.3) using ggtree (v3.10.1) (Xu *et al*., 2022), ggplot2 (v4.0.0) (Wickham, 2016), ape (v5.8-1) (Paradis & Schliep, 2019), EnhancedVolcano (v1.20.0) (Blighe, 2026), pracma (v.2.4.4) (Borchers, 2025).

### Pathogenicity assays

Pathogenicity on cherry (scion: Coral Champagne, rootstock: Gisela 6) was assessed using assays from (Hulin *et al*., 2018b). Syringe-infiltration assays were performed on 2-week-old detached leaves with an inoculum of 1 x 10^6^ CFU/mL in 10mM MgCl_2._ Leaves were placed in containers with 15% water agar overlaid with paper towel and sealed using plastic bags to generate high humidity. Leaves were incubated in a growth chamber at 24°C, 16h light : 8h dark. Bacterial populations were assessed via dilution plating on KBA with rifampicin and cycloheximide. Woody tissue inoculations were performed on dormant 1-year old trees using an inoculum of 1 x 10^8^ CFU/mL in 10mM MgCl_2._ Three separate biological replicates per strain with six inoculations per tree were performed. Trees were stored at 20°C, 16h light: 8h dark for 10 weeks. Lesion area was calculated against the reference scale in ImageJ (Schindelin *et al*., 2015).

### Phenotypic and growth assays

Biofilm formation was performed as in (Peng *et al*., 2020) using full strength LB media and etched 96 well plates as in (Davies & Marques, 2009). For motility assays, methods from (Ichinose *et al*., 2013) were used. For growth curves, overnight KB cultures were centrifuged to pellet cells and pellets washed and resuspended to an OD_600_ of 0.02 in KB or LB HIM in 96 well plates and OD monitored in a TECAN Infinite M Plex plate reader. As Pam failed to grow in HIM in the plate reader, a larger culture (100 mL) was grown from OD_600_ 0.1 for 72 h.

### Creation of Pam mutants

Pam mutants were created as described in (Kvitko *et al*., 2007) with pK18msB (Ling *et al*., 2022) (Addgene #177839) using ClonExpress Ultra One-Step Cloning Kit (Vazyme). Constructs were confirmed by sequencing (Plasmidsaurus). Constructs were mated into Pam using triparental mating as in (Neale *et al*., 2020). Selection of transconjugant Pam was performed on KBA with kanamycin, rifampicin and nitrofurantoin. Sucrose counter-selection was performed on LBA (without salt) with 15% (W/V) sucrose. Resulting colonies were assessed for deletion by colony PCR. For quantifying protein expression, genes with added C-terminal HA tags expressed under their native promoter were synthesized (Twist BioScience) and cloned into pUCP20-TK (Zhou *et al*., 2009) orientated in the reverse direction to the lac promoter. Plasmids were transformed into Pam via electroporation as in (Choi *et al*., 2006), except recovery was for 2 h at 28°C.

### qPCR analysis of bacterial and plant gene expression during infection

qPCR was used to assess expression of plant and bacterial genes during infection. Bacteria inoculum at OD_600_ of 0.5 (∼2.5 x 10^8^ CFU/mL) was infiltrated into detached leaves. Each strain was inoculated 30 times onto each leaf (∼100mg of tissue). After 6 or 24hr, leaf discs were frozen in liquid nitrogen and stored at −80°C. RNA was extracted from 100 mg of tissue using the FastPure Universal Plant Total RNA isolation kit (Vazyme) and subsequently treated with Turbo DNA-Free Kit (Thermo Fisher Scientific). 500ng RNA was converted to cDNA using superscript III (Thermo Fisher Scientific) with either random hexamers or oligoDT_20_ for bacterial and plant gene expression respectively. For plant genes, the *Prunus avium* genome (NCBI assembly GCF_002207925.1) was used for qPCR primer design. For bacterial genes, the IR 634 genome (GCF_054719885.1) was used.

qPCR was performed using the CFX opus 96 (Bio-Rad) 3-step amp and melt protocol. All reactions were performed using SsoAdvanced SYBR Green Supermix (Bio-Rad) with primers at 0.2µM and 3.125ng total cDNA. RT-qPCR reactions were run in sets of biological replicates with three technical replicates per sample. Runs were standardized using amplification of Pam IR 634 *rpoD* from 40ng gDNA. Expression was normalized against Elongation Factor alpha (LOC110771043) for plant gene expression and *rpoD* (ACYHMY_24945) for bacterial gene expression.

### Immunoblotting

Pam expressing HA-tagged genes were grown overnight in KB, adjusted to OD_600_ 0.4 and centrifuged at max speed for 5 minutes. The pellet was washed twice in 10 mM MgCl_2_ and resuspended in HIM to an OD_600_ 0.4 and incubated shaking at 17°C to induce T3SS expression. After 24h, 10mL cultures were standardized to OD_600_ 0.4, centrifuged at 13,000 RPM for 5 min and pellets resuspended in 100 µL of Laemmli sample buffer (Fisher scientific) and boiled for 5 min at 95 °C. Immunoblotting was performed as in (Zhou *et al*., 2009), except blocking used 5% dehydrated milk powder, followed by incubation with anti-HA antibody, mouse monoclonal (Millipore Sigma) at 1:5000 and Goat Anti-Mouse IgG Antibody HRP conjugate (Millipore Sigma) at 1:10000. To quantify protein expression, band intensities were measured using pixel density in ImageJ with the background density used as a blank.

## Results

### Prophages are widespread across the *P. syringae* species complex

Our previous work revealed that the T3E gene *hopAR1* is carried on a prophage PamPP1 in Pam (Hulin *et al*., 2023b). To assess the prevalence of prophages across the PSSC, we predicted 4107 prophages across 1669 genomes (Figure 1A). Of these, 3638 were predicted as intact and contained a “head” gene found within 1176 genomes. This indicates that 70% of strains possess potentially active prophages (Table S3). To define taxonomic groups, phage sequences were run through VipTreeGen and vConTACT3 (Nishimura *et al*., 2017; Bolduc *et al*., 2025). The PamPP1 prophage belonged to VipTree group 26 and was part of a novel family (12) within a novel order (45) within the Caudoviricetes (Figure 1A, full data for all prophages are in Figure S1). VipTree group 26 contained 620 prophages and was distributed across the PSSC (Figure 1B). 454 of these prophages were predicted intact indicating likely functionality. A lack of clustering of phage sequences by phylogroup suggests this group transferred frequently across the PSSC. 17 additional temperate phages from the Virus-Host database were within this group (Table S3) including phages infecting other *Pseudomonas* species such as *P. fluorescens*, *P. aeruginosa* and *P. putida,* indicating related phages are found outside *P. syringae*.

**Figure 1:**
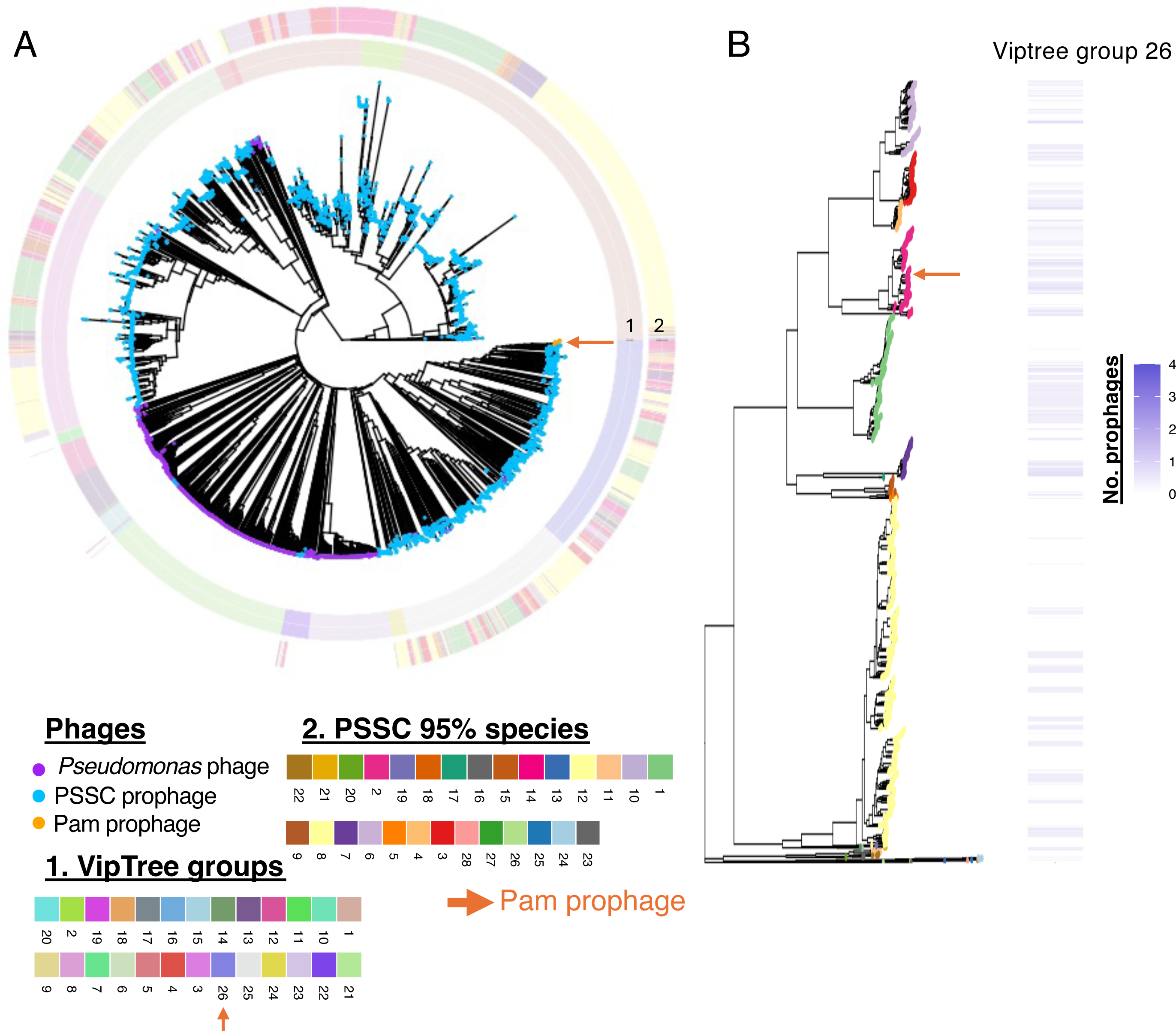
The PSSC contain s divers eprophages and the prophages related to the PamPP1 prophage are found across the complex. (A) VipTree phylogeny of PSSC prophages with other *Pseudomonas* phages belonging to the Caudoviricetes. PSSC prophages are highlighted with blue circles and *Pseudomonas* phage with purple. The PamPP1 prophage is highlighted in orange and with an arrow. The first circle on the heatmap shows VipTree groups identified using hierarchical clustering, whilst the second circle shows which *P. syringae* species (based on 95% average nucleotide identity) each prophage was identified within. (B) A core genome phylogenetic tree of 1669 strains from across the PSSC. Tips are highlighted by *P. syringae* species at the 95% average nucleotide identity level (colors the same as in A). The heatmap shows the presence of VipTree phage group 26 across the PSSC.

### Conservation of *hopAR1-*carrying prophage in *Pseudomonas amygdali*

To assess the conservation of the PamPP1 prophage within *P. amygdali*, we performed a focused search for prophages within this sub-phylogroup (Figure 2A). All Pam strains possessed *hopAR1*, as did pv. *myricae* and all but one strain of pv. *eriobotryae*. All genomic regions carrying *hopAR1* had phage genes present, however for pv. *myricae* prophages were incomplete, lacking genes for particle production while pv. *eribotryae* prophages were both predicted as complete and incomplete. Alignments of the region showed that the PamPP1 prophage is conserved in Pam yet fragmented in other pathovars (Figure S2). Prophages within pv. *myricae* only retained lysis genes downstream of *hopAR1* and a region containing a gene encoding a predicted Toll-Interleukin-1 Receptor protein. However, pv. *eribotryae* prophages maintained the lysis genes and one of the tail proteins with 100% homology to those in Pam, while other predicted phage structural proteins showed 0-50% homology. The annotated Pam prophage (Figure 2B) showed most genes encoded protein functions related to phage assembly and bacterial lysis (Table S4). A major category of predicted bacterial (moron) proteins was transcriptional regulators (13 genes). Other functions included a protein structurally homologous to an invasion-associated protein B (PDB: 3DTD) from *Bartonella henselae* which could have a role in pathogenicity, a pseudogenized T3E HopBK1, defense proteins involved in bacterial immunity and enzymes (such as hydrolases).

**Figure 2:**
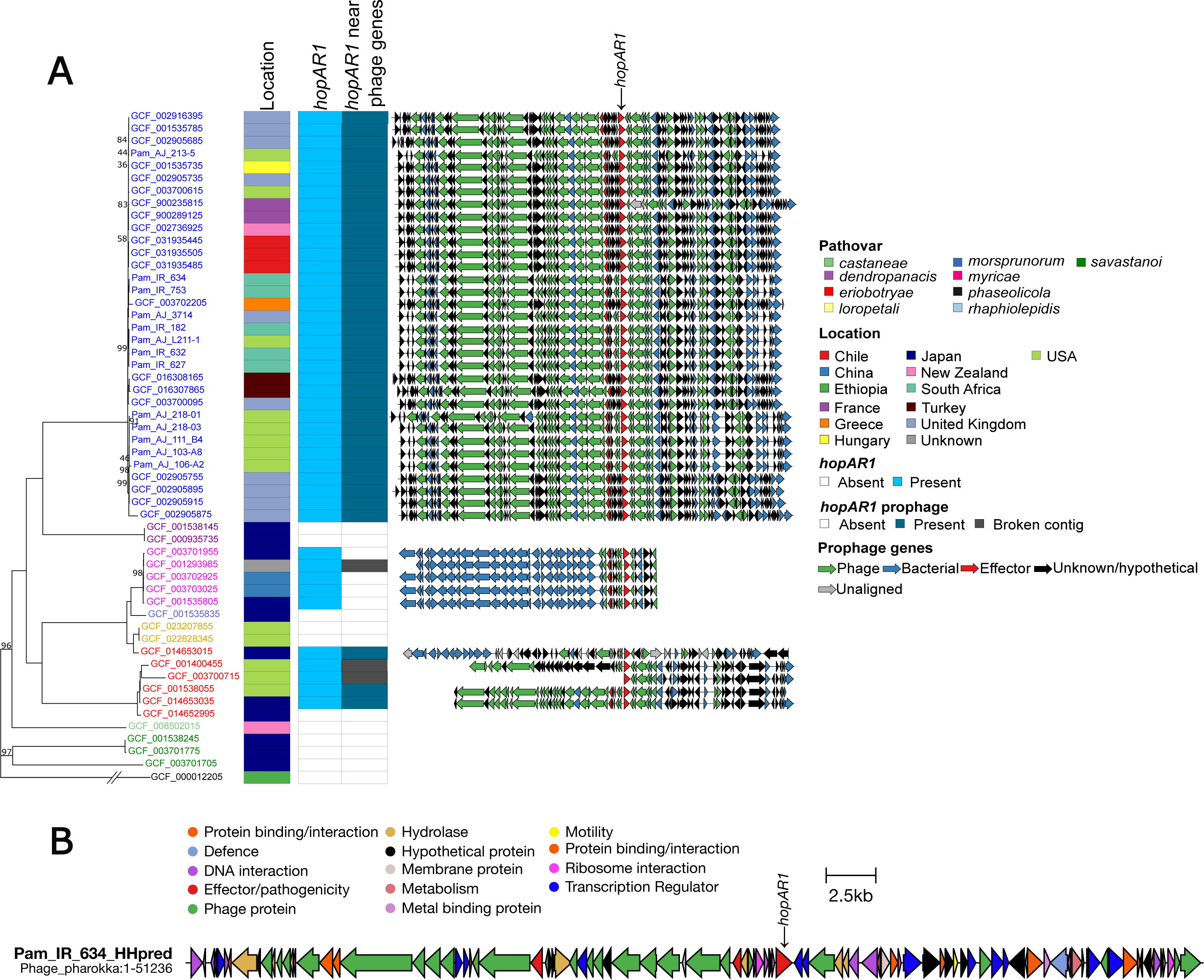
The PamPP1 prophage is highly conserved within Pam and is present in other pathovars based on sequence homology. The region contains genes with a range of functions that could impact the bacterial host. (A) Phylogeny of phylogroup 3 *P. amygdali* based on the core genome alignment for publicly available genomes. The tree was rooted using *P. savastanoi* pv. *phaseolicola* 1448A as the outgroup. Bootstrap values <100 are shown at branch points. Overall, nine pathovars were included and can be identified by the colored accession numbers/names in the tree. The heatmap shows location of isolation, the presence of *hopAR1* as detected by HMMER analysis, and the prediction of a prophage in that region via PHASTEST in column three. Prophages aligned via clinker are presented next to each strain with genes colored by function, and unaligned genes to the closest neighbor are shown in grey. (b) The PamPP1 prophage region in strain IR 634 with functional annotations predicted with HHPred. Genes predicted to play a role in phage particle assembly and functions such as lysis etc. are shown in green (predicted via PHASTEST) while other predicted functions are shown with other colors.

### Deletion of the prophage region causes altered phenotypes not explained by the loss of *hopAR1*

To examine the importance of the conserved T3E *hopAR1* and the PamPP1 prophage in Pam, deletion mutants of the effector and whole prophage were created for the pathogenic strain IR 634 (IR 634:Δ*hopAR1*, IR 634:ΔP). Deletion of the 51 Kb prophage region significantly reduced virulence within leaf tissue, with visible differences in lesion formation (Figure 3A) and reduced ability to grow within the leaf, as populations at 7 days post inoculation were over 1-log lower than the wildtype (p=0.009). Likewise, IR 634:ΔP showed a significant reduction in lesion severity and size in woody tissue (p=<0.001) (Figure 3B). The wildtype caused typical black necrosis and gumming, whilst IR 634:ΔP caused brown lesions with reduced gumming. Meanwhile IR 634:Δ*hopAR1* showed no significant reduction in virulence in detached leaf assays compared to the wildtype (Figure S3).

**Figure 3:**
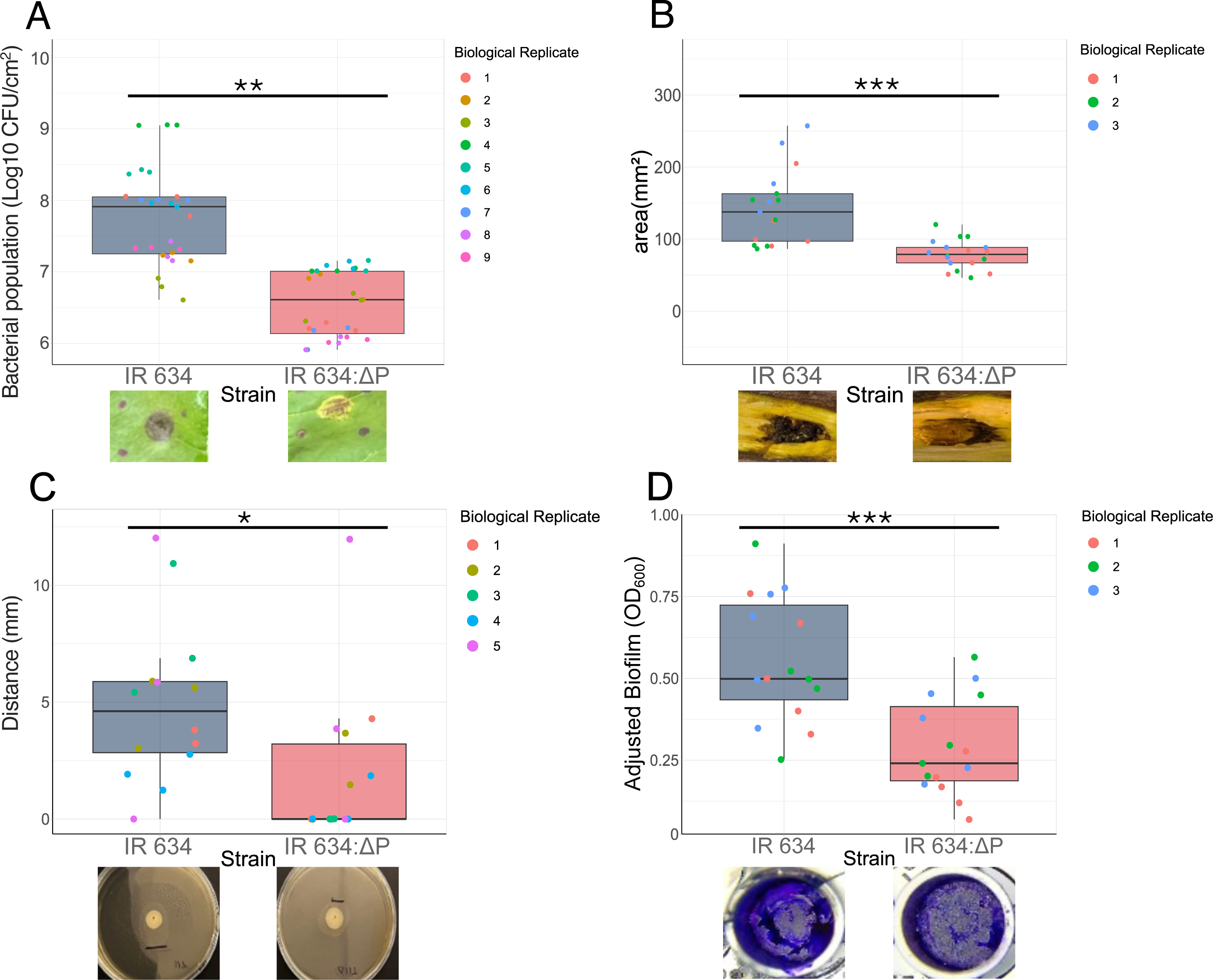
Deletion of the PamPP1 prophage in Pam IR 634 leads to altered phenotypes. (A) Bacterial population count assays comparing populations of the prophage deletion scarless mutant IR 634:ΔP to the wildtype strain IR 634 in leaf tissue 7 days post inoculation. Below are representative images of the lesions caused in leaves when infiltrated with their respective strain. (B) Lesion area two months after inoculation of woody tissues. Below are representative images of the lesions in the experiment. (C) Twitching motility on 1 % KBA after 48 h and an example image of IR 634 and IR 634:ΔP where visible differences between the distance moved are marked. (D) Biofilm formation in a 96-well plate following 48 h inoculation, staining and dilution. Representative images of the adhered mature biofilm after staining and washing can be seen. For all graphs, significant differences were tested via Welch two-sample t test (Significance (P<0.05 =*, P < 0.01=**, P < 0.001= ***).

IR 634:ΔP and wildtype were then assessed under laboratory conditions for motility and biofilm formation. While the strains were not observed to swim or swarm consistently in laboratory conditions (data not shown), twitching was reduced in IR 634:ΔP compared to wildtype (p=0.03) (Figure 3C). Likewise, mature biofilm formation was significantly reduced in IR 634:ΔP (p=<0.001) (Figure 3D). To determine if the phenotypic differences observed were due to defective growth of the mutant, *in vitro* growth was assessed in rich media KB and LB and minimal medium HIM (Figure S4). Overall, no significant difference was observed in the growth between the wildtype and IR 634:ΔP in these media.

### The Pam prophage rewires transcription during *in vitro* growth affecting expression of the T3SS

We hypothesized that phenotypic differences between the wildtype and IR 634:ΔP could be due to transcriptional changes driven by prophage-encoded regulators. RNA-Seq was conducted in KB and after transfer to HIM (0, 5 and 25 h). Principal Coordinate Analysis revealed clear separation of mutant and wildtype transcriptomes, with the largest divergence at 5 h and 25 h (Figure S5). Heatmaps of all time points are shown in Figure S6 and sequencing statistics are in Table S5. Normalized read counts and DEGs are in Table S6. Specific genes of interest are in Table S7.

In KB (Figure S7), minimal transcriptional differences were observed, with only five genes outside the prophage region differentially expressed, primarily involving downregulation of motility and chemotaxis functions. Upon transfer to HIM (0 h), minor differences were seen (Figure 4A). The largest response (combined Log2FC and p-value) was in the downregulation of an IS91 transposon in IR 634:ΔP, while the largest upregulation observed was in *hrpQ* (*sctD*) which encodes a T3SS component. Upregulation of the T3SS pathway was evident in IR 634:ΔP, suggesting more rapid activation compared to the wildtype. By 5 h in HIM, major transcriptional differences were observed. IR 634:ΔP showed widespread downregulation across multiple pathways compared to wildtype, including alginic acid metabolism, general metabolism (glycolysis, pentose phosphate), starch and sucrose metabolism, and osmotic stress responses (Figure 4B). T3SS genes remained higher in IR 634:ΔP, indicating sustained activation. At 25 h, the greatest divergence between strains was observed (Figure 4C). IR 634:ΔP exhibited strong upregulation of translation-related pathways, including ribosomal proteins, alongside upregulation of 16 T3SS structural genes. In contrast, few pathways were downregulated, indicating a shift toward enhanced protein synthesis and virulence-associated functions.

**Figure 4:**
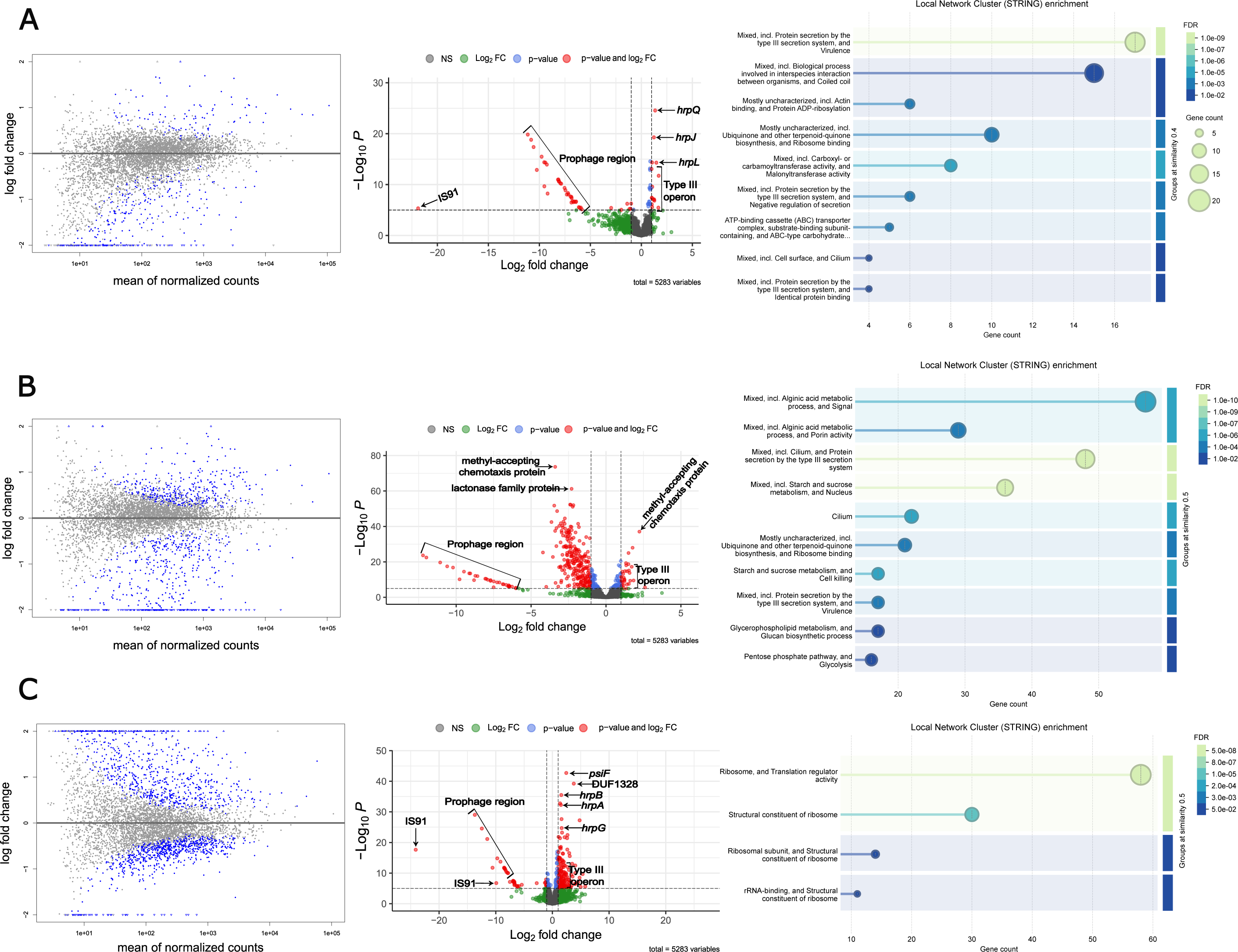
Deletion of the PamPP1 prophage in Pam leads to significantly altered gene expression at 0, 5 and 25 h in HIM between IR 634 and IR 634:ΔP. (A) Is analysis at 0 h, with 264 genes with an adjp < 0.05. (B) is analysis at 5h with 745 genes with an adjp < 0.05. (C) is analysis at 25h with 1241 genes with an adjp < 0.05. Each analysis shows the MA plot (Left) showing the differential expression against the expression level of all genes at the same time point of IR 634:ΔP against IR 634, genes with a padj < 0.05 are shown in blue. The enhancedvolcano plot analysis (middle) in which genes with a log2FC >2 are highlighted in green and denoted by vertical dashed lines, a p-value <10e-6 are shown in blue and cut-off denoted by horizontal dashed lines, genes displaying both those values are shown in red and non-significant genes are shown in grey. Significantly differentially expressed genes of interest are labeled based on their locus tag. The Local Network Cluster (Right) identified through STRING analysis against the proteome of Pam IR 634 (STRG0A01MPX) terms are sorted by the gene counts and then grouped with a similarity of ≥ 0.8 and merged with a similarity of ≥ 0.6 based on the Jaccard index of the gene sets within terms according to their p-values to prioritize the strongest statistical significance of all gene with a padj < 0.05.

Most differentially expressed genes were time-specific (Figure S8) and predominantly downregulated in the mutant. In contrast, the T3SS operon was consistently upregulated in IR 634:ΔP. We examined genes involved in T3SS assembly and effectors (Table S8) in more detail. Most striking was the assembly genes which are encoded in the *hrp*/*hrc* pathogenicity island including multiple operons (Collmer *et al*., 2000) (Figure 5A), where almost all genes exhibited higher expression in IR 634:ΔP. The RNA polymerase sigma factor *hrpL* was higher in IR 634:ΔP at both 5h and 25 h. HrpL controls T3SS operon expression and its higher expression could lead to upregulation of other T3SS genes. T3E genes shown in Figure 5B were variable, with some genes expressing higher in the wildtype and others in IR 634:ΔP. Many effectors were expressed at a higher amount in IR 634:ΔP compared to wildtype by 5h and remained significantly higher at 25 h.

**Figure 5:**
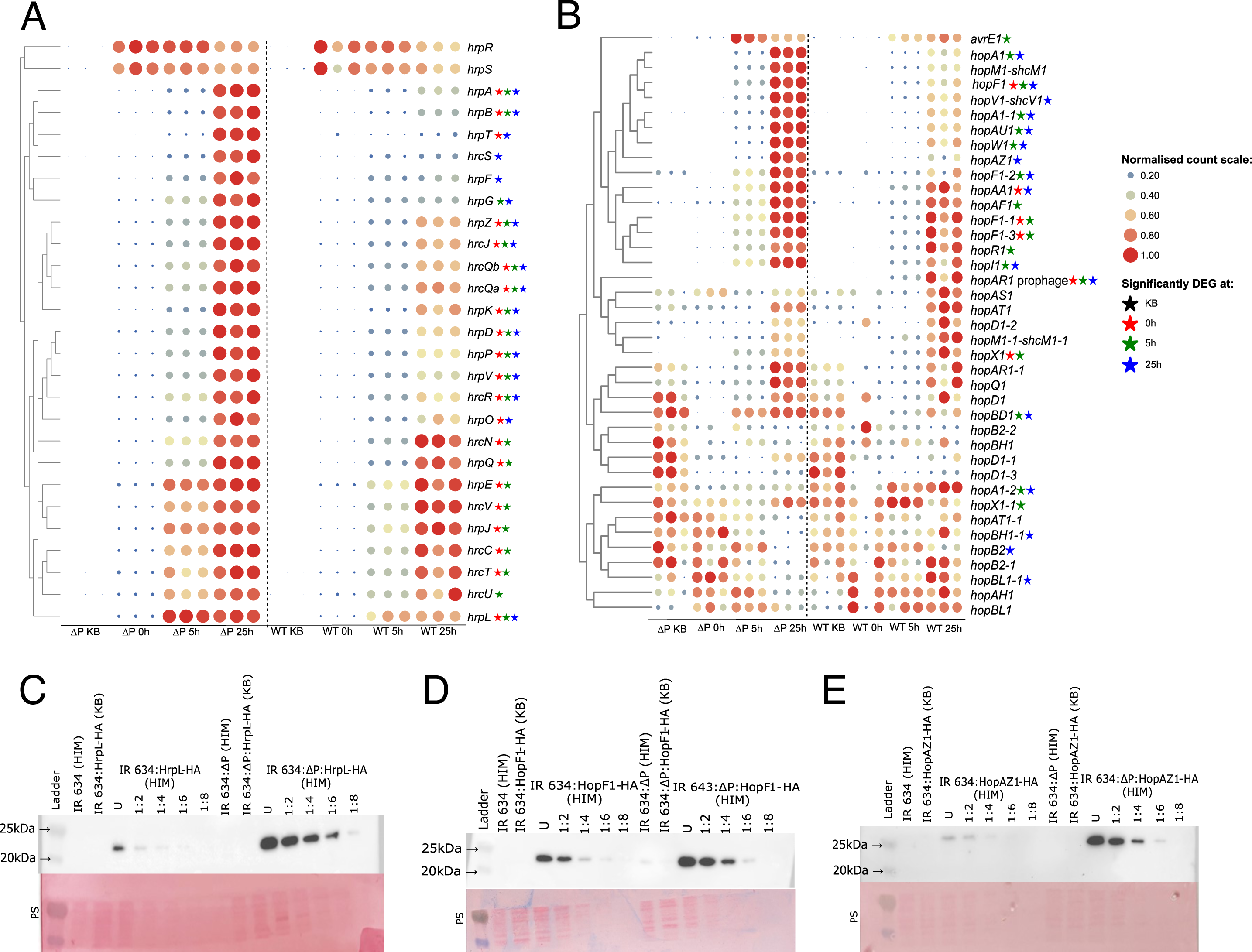
The PamPP1 prophage alters expression of T3SS and T3E genes in Pam. Analysis of the T3SS and associated T3E expression in IR 634 and IR 634:ΔP using normalized RNA sequencing read counts and quantifying HA-tagged proteins via western blotting. (A) Analysis of the T3SS operon in IR 634 (ACYHMY-07265 to ACYHMY-07395). (B) Analysis of all T3SS effectors identified in IR 634 genome by HMMER analysis against the *Pseudomonas syringae* Type III Effector Compendium database with a bitscore > 50. (A) and (B) were made using the normalized read counts from DEseq2 analysis, data is displayed as a normalized count scaled by row of 0 to 1, then clustered based on expression patterns. Significance at different time points is signified through colored stars. (C) HA-tagged HrpL protein expression at 24h. (D) HA-tagged HopF1 expression at 24 h. (E) HA-tagged HopAZ1 at 24 h. All western blots were performed under the same conditions. From left to right is the ladder, IR 634 grown in HIM, IR 634 carrying the HA-tagged gene grown in non-inducing media (KB), IR 634 carrying the tagged gene grown in HIM, with a serial dilution series from undiluted (U) to 1:8 dilution, followed by IR 634:ΔP grown in HIM, IR 634:ΔP carrying the HA-tagged gene grown in non-inducing media (KB), IR 634:ΔP carrying the tagged gene grown in HIM, with a serial dilution series from undiluted (U) to 1:8 dilution to allow for standard curve quantification. Below each plot is the complementary ponceau stained (PS) membrane.

Overall, expression of T3SS-related genes was significantly altered in IR 634:ΔP compared to the wildtype. This could be due to altered expression of known regulators of the T3SS (Xie *et al*., 2019) (Figure S9A, Table S7). The master upstream regulators *hrpR* and *hrpS* did not show differential expression. Whilst, regulators such as *algU*, four homologs of *chp8*, *relA*, *rhpC* and *rhpP* were all downregulated at certain times in HIM in IR 634:ΔP, whilst the regulator *spoT* was upregulated in IR 634:ΔP compared to wildtype. To assess if T3SS expression differences observed in the transcriptome were also seen at the protein level, western blots of HA-tagged HrpL and T3E genes HopAZ1 and hopF1-3 were performed and protein level quantified (Figure 5C-E, graphs in Figure S10). This confirmed that after 24 h of growth in HIM, IR 634:ΔP produced these proteins at a higher level than the wildtype.

To see if the transcriptome data could explain other altered phenotypes observed, we also examined motility associated genes (flagella and taxis). Expression patterns of flagellar genes (Figure S9B) did not show a clear pattern across the two strains, many of the assembly proteins were upregulated at 5 hours in IR 634:ΔP but downregulated compared to wildtype at 25 hours. In terms of twitching, the *pilB* associated ATPase (Figure S9C) was more strongly expressed in the wildtype at 25 h compared to IR 634:ΔP. Chemotaxis associated genes likewise did not show large differences in their expression, although some were downregulated in IR 634:ΔP in HIM compared to wildtype (Figure S9D).

### Deletion of the prophage leads to mis-regulation of the T3SS and T3Es *in planta* and differences in expression of plant immune responses

The *in vitro* transcriptome data indicated that T3SS expression was reduced in the wildtype indicating the presence of the PamPP1 prophage directly or indirectly lowers its expression in HIM. To assess if these differences were also seen *in planta* and examine how this could relate to identified virulence phenotypes, cherry leaves were inoculated with wildtype and IR 634:ΔP and qPCR performed at 6 h and 24 h to examine T3SS gene expression (Figure 6A-F). At 6 h, expression of all 6 of the T3SS genes screened was significantly higher in the wildtype than in IR 634:ΔP. However, at 24h the inverse was seen where IR 634:ΔP expressed four of the observed T3SS genes at a significantly higher level than the wildtype.

**Figure 6:**
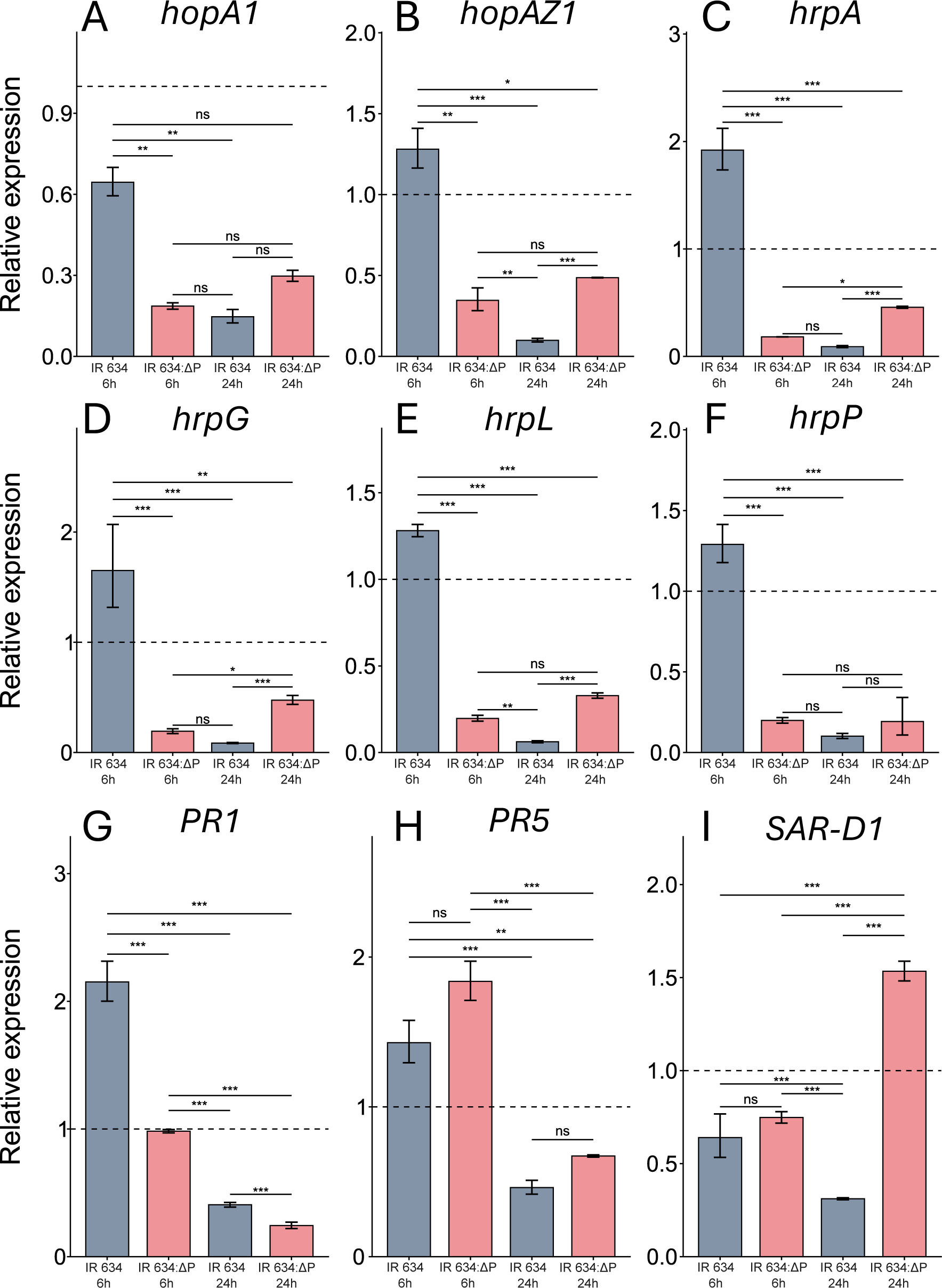
Relative expression of the T3SS, T3 effectors in Pam IR 634 and IR 634:ΔP and host cherry immune response genes at 6h and 24h during infection of leaves. Graphs are labelled by the gene analyzed. All graphs show a dashed line indicating a relative expression of 1. Significance (P<0.05 =*, P < 0.01=**, P < 0.001= ***) is denoted by stars above according to ANOVA with Shapiro-Wilk Normality test. Data is the combination of biological replicates (n=3) and three technical replicates (n=3) with standard error of the mean. For the effectors *hopA1* (ACYHMY_07405) and *hopAZ1* (ACYHMY_09705) and the type three regulatory components *hrpA* (ACYHMY_07275), *hrpG* (ACYHMY_07310), *hrpL* (ACYHMY_07390) and *hrpP* (ACYHMY_07360), relative expression was calculated using the ΔΔCq value against the housekeeping gene *rpoD* (ACYHMY_24945). For cherry immune response genes pathogenesis-related protein 1-like (PR1, LOC110769299), pathogenesis-related protein 5 (PR5, LOC110762616) and systematic acquired resistance deficient like protein 1 (SAR-D1, LOC110754122), relative expression was calculated using the ΔΔCq value against the housekeeping gene Elongation-factor α-like (EF-α, LOC110771043).

The T3SS is key to delivering effector proteins into plant cells to suppress immunity. We hypothesized that differences in T3SS and T3E expression could lead to differences in how the plant immune system responds during infection of these strains. We tested the expression of three putative plant immunity genes which are induced during immune responses towards *P. syringae* in cherry (Lienqueo *et al*., 2024). This showed their expression differed during responses to the wildtype and IR 634:ΔP as shown in Figure 6G-I. At 6 h the wildtype appears to significantly trigger higher expression of the PR1-like gene in comparison to IR 634:ΔP. The systematic acquired resistance deficient like protein 1 (SAR-D1) gene appears to show no difference between the strains at this point. However, leaves inoculated with IR 634:ΔP did show a slight, although not significant upregulation of the PR5-like gene compared to wildtype. By 24 h, a larger difference can be seen in which SAR-D1 gene expression was over three-fold higher in leaves inoculated with IR 634:ΔP versus those inoculated with the wildtype. Interestingly, the PR5-like gene was also more highly expressed in IR 634:ΔP-inoculated leaves compared to the wildtype, though not significantly.

### Deletion of *hrpL* returns strains to the same level of virulence *in planta*

Our results suggested the PamPP1 prophage alters bacterial transcriptome, affecting genes involved in bacterial responses to the environment, metabolism and virulence gene expression. The reduced virulence in planta could be due to various altered phenotypes. To assess if the reduced virulence of IR 634:ΔP *in planta* was specifically linked to altered T3SS expression, rather than altered survival due to other phenotypes, *hrpL* knockout mutants were generated for both IR 634 and IR 634:ΔP. While the reduced virulence of IR 634:ΔP was consistent, removal of *hrpL* made both strains non-pathogenic. Populations were found at the same level *in planta* indicating that the virulence differences observed could be dependent on T3SS differences rather than altered general survival mechanisms where we might expect IR 634:ΔP to still be reduced compared to wildtype (Figure 7).

**Figure 7:**
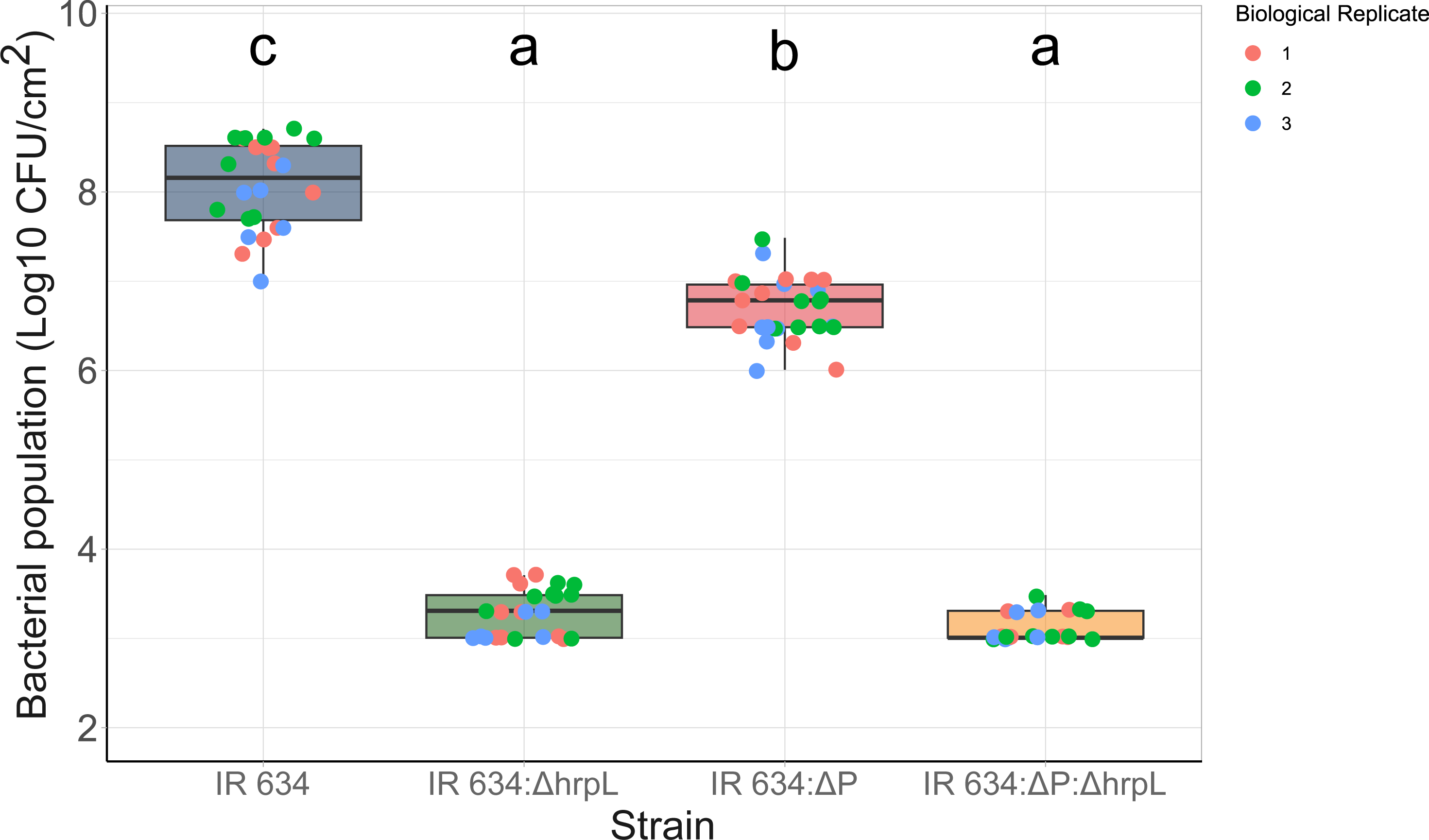
Removal of the T3SS regulator *hrpL* causes IR 634 and IR 634:ΔP to persist at the same level *in planta.* Bacterial population count assays comparing populations of IR 634:ΔP, the double knockout IR 634:ΔP:Δ*hrpL* and IR 634:Δ*hrpL* mutant to the wildtype strain IR 634 in leaf tissue. Significant differences tested via ANOVA (P<0.05 Tukey post-hoc testing) are shown by the compact letter display groupings above.

## Discussion

In this study we determined that a prevalent prophage PamPP1 was required for full virulence of the cherry pathogen Pam, influencing several phenotypes such as the T3SS expression, biofilm and type IV twitching motility. Integrated prophages are widespread across bacteria and enriched in pathogens (Inglis *et al*., 2025). We found that ∼70% of PSSC genomes contain intact prophages, suggesting potential functionality. Our previous work identified PamPP1 can be transferred between strains (Hulin *et al*., 2023b). This prophage belongs to a Caudoviricetes family distributed across the PSSC which has been frequently gained and lost across phylogroups, indicating a role in shaping evolution and ecology of these plant pathogens.

Analysis of the PamPP1 prophage in phylogroup 3 showed conservation, with homologous phage sequences in related pathovars (*myricae* and *eriobotryae*), suggesting it was acquired prior to divergence. Although the T3E *hopAR1* was conserved, its deletion did not significantly alter virulence in detached leaf assays, consistent with studies showing limited phenotypic effects of individual T3E deletions (Jayaraman *et al*., 2023; Hemara *et al*., 2025). In contrast, deletion of the entire prophage reduced virulence and impaired biofilm formation and type IV pili twitching. These findings suggest the prophage contributes broadly to bacterial physiology and may have facilitated the emergence of these pathogens (Figure 8).

**Figure 8:**
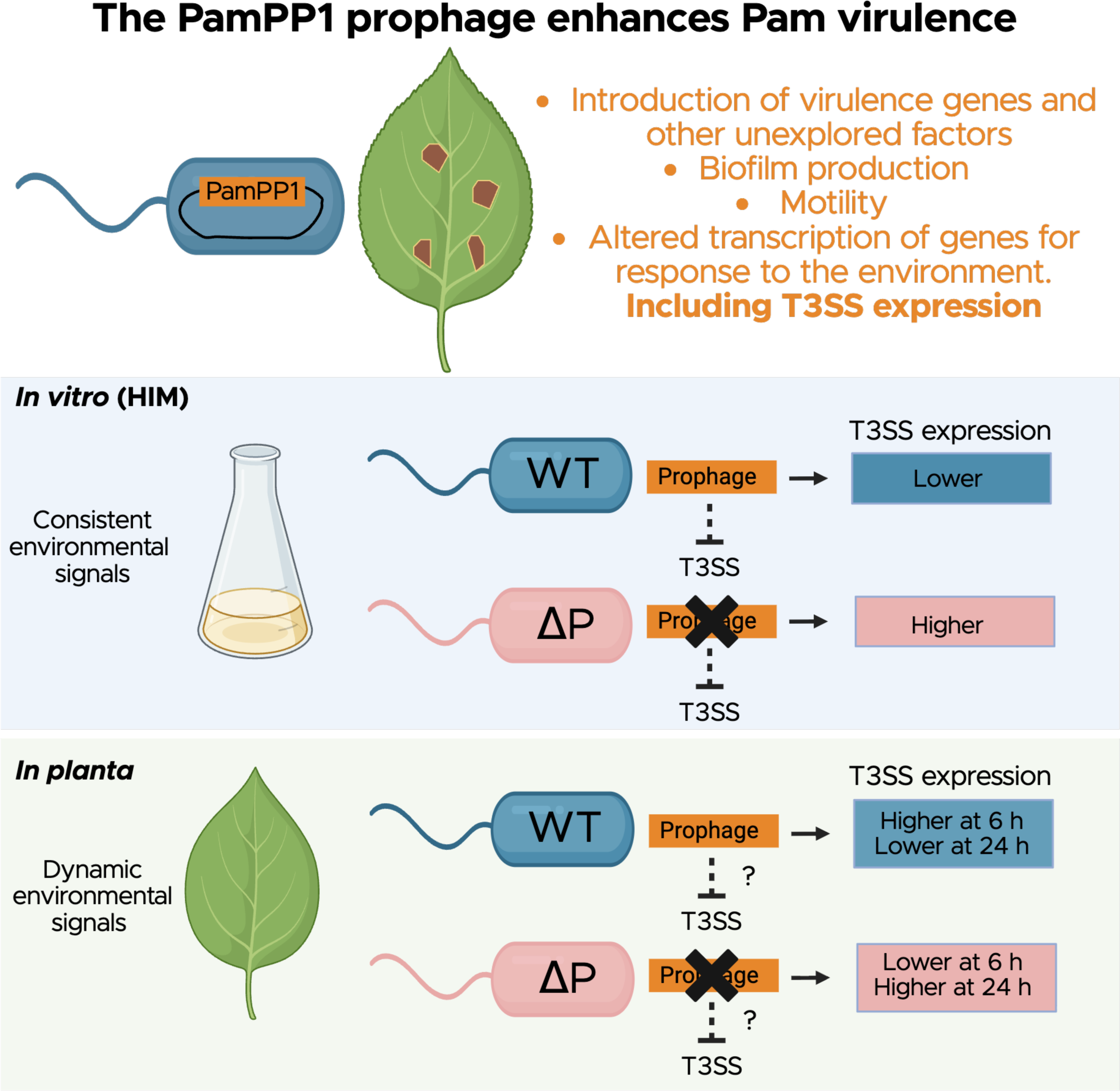
Working model of how the PamPP1 prophage contributes to Pam virulence with emphasis on our findings that T3SS and T3E gene expression is altered by the presence of the prophage.

Using a variety of functional annotation tools, we predicted the prophage contains genes encoding 13 transcriptional regulators. Given this, RNA-seq was used to examine gene expression over 25 h in HIM, which mimics conditions experienced in the plant during infection (Huynh *et al*., 1989). The wildtype and IR 634:ΔP showed substantial divergence in gene expression, particularly in T3SS pathways. Prophages are known to regulate host genes, for example the phage transcription factor Cro can activate T3SS expression in *Escherichia coli* (Hernandez-Doria & Sperandio, 2018).

In rich KB medium, only five genes outside the prophage were differentially expressed, indicating minimal impact in these conditions. However, divergence increased in HIM, suggesting the prophage modulates transcriptional responses in this environment. At 0 h, actin-binding genes were downregulated in IR 634:ΔP. Actin is crucial for bacterial movement and T3SS assembly (Stevens *et al*., 2006; Russo *et al*., 2021). At 5 h, genes involved in alginic acid metabolism and regulation were strongly affected. The AlgU sigma factor regulates stress responses, motility, and T3SS expression in *P. syringae* pv. *tomato* DC3000 (Markel *et al*., 2016), while AlgR contributes to pathogenicity and biofilm formation (Peñaloza-Vázquez *et al*., 2004; Huang *et al*., 2022). Downregulation of AlgR and 46 of 57 alginate pathway genes in IR 634:ΔP suggests the phage upregulates this key network. Similarly, reduced expression of starch and sucrose metabolism genes, which have been linked to virulence in *P. syringae* pv. *Actinidiae* (Asif *et al*., 2025), indicates broader metabolic mis-regulation in the mutant. By 25 h, enrichment of ribosome and protein export genes among DEGs suggests altered translational capacity.

The most striking effect was on T3SS and T3E expression. *In vitro*, these genes were more highly expressed in IR 634:ΔP indicating repression by the prophage. Protein-level differences at 24 h confirmed this pattern. *In planta*, however, dynamics differed: at 6 hours post inoculation, several T3SS genes were upregulated in the wildtype, whereas by 24 hours IR 634:ΔP showed higher expression. This discrepancy likely reflects differences between HIM and the plant apoplast. The plant environment includes host-derived signals, nutrients, and immune responses which are absent *in vitro* (Haapalainen *et al*., 2009; O’Malley *et al*., 2023). Deletion of *hrpL* abolished virulence differences, indicating that altered T3SS regulation may underlie differences in virulence rather than altered general survival.

The T3SS and T3Es are key virulence factors that suppress host immunity, but their effectiveness depends on precise timing and dosage. Because they are energetically costly, their expression must be tightly controlled (Xie *et al*., 2019). Prolonged activity may reduce growth; however, no growth differences were observed between the wildtype and IR 634:ΔP in HIM, indicating that reduced virulence is unlikely due to general growth differences. Instead, we propose that mis-timed T3SS expression leads to ineffective immune suppression or inappropriate immune activation. *In planta*, the wildtype rapidly induced and then downregulated T3SS expression by 24 h, whereas IR 634:ΔP showed delayed induction and prolonged expression. Although essential for virulence, excessive or prolonged T3SS activity can trigger host immunity and reduce pathogenicity (Tampakaki & Panopoulos, 2000; Waite *et al*., 2017; Laflamme *et al*., 2020; Jin *et al*., 2023). Consistent with this, T3SS expression differences correlated with altered host immune responses: PR1 was upregulated in wildtype infections at 6 hours, while SAR-D1 was strongly induced in IR 634:ΔP infections at 24 h. This suggests distinct host responses contribute to differences in virulence. Further transcriptomic analysis of the plant during infection would clarify broader immune dynamics.

Timing of T3SS expression varies across plant pathogens. *Dickeya dadantii* on potato peaks at 12 h and declines by 48-72 h (Cui *et al*., 2018), whereas *Ralstonia solanacearum* maintains expression throughout tomato infection (Monteiro *et al*., 2012). In *Pseudomonas savastanoi* pathovars *savastanoi, nerii* and *fraxini*, T3SS gene expression in HIM peaked between 3-18 h and declined by 24 h (Tegli *et al*., 2011), while in *P. savastanoi* pv. *phaseolicola* expression in bean plants peaks at 2-4 h and is absent by 24 h (Thwaites *et al*., 2004). The expression of the T3SS is also heterogeneous at the single-cell level (López-Pagán *et al*., 2024), coordinated by environmental cues (Caullireau *et al*., 2025). Tight regulation is required to balance immune suppression with avoidance of detection, particularly during transitions from biotrophic to necrotrophic phases. Small timing shifts may therefore alter infection outcomes (Xie *et al*., 2019). This is supported by the identification of natural mutants of regulatory genes that exhibit altered virulence (Xie *et al*., 2023) Our results suggest prophages contribute to this regulatory complexity. We hypothesize alterations in the expression of putative regulators on the prophage could be behind differences seen in HIM and *in planta*. As mobile elements, prophage gene expression could depend on their state (integrated or excised) and stage in their lifecycle (lysogenic or lytic). These transitions are intrinsically linked to bacterial stress and thus could change *in planta* during infection with downstream consequences for virulence regulation. Prophage genes were found to be upregulated in *P. syringae* pv. *syringae* in the apoplast during infection of bean (Yu *et al*., 2013). Further examination of PamPP1 gene expression during infection would be required to examine this hypothesis.

Overall, prophages are widespread in the PSSC and represent an important component of the accessory genome influencing plant-associated fitness (Holtappels *et al*., 2024). Their distribution suggests these findings extend beyond Pam. The removal of PamPP1 in Pam caused widespread transcriptional changes affecting virulence and motility, with a major impact on T3SS regulation, particularly *in planta*. This indicates prophages play a broader role than just carriage of virulence factors. Future work should identify the specific prophage factors responsible for directly or indirectly altering T3SS expression. As expression of canonical regulators such as HrpR/S were unaffected other pathways may be involved (Xie *et al*., 2019). Our findings highlight prophages as important drivers of bacterial evolution. Further understanding these phages, how they persist and their bacterial host range within in epiphytic communities will offer insight and opportunity to predict disease emergence.

## Acknowledgements

We thank Michigan State University staff Chrislyn Patricka, Rachel Wood, Cody Keilen and Cory Outwater for advice on plant growth conditions. We thank Dr Xufeng Wang for advice on visualizing RNA-Seq analysis. We are grateful to Dr John Mansfield and Dr Amelia Lovelace and the rest of the Hulin lab for their critical reading of the manuscript.

## Author contributions

D.M: Conceptualization, investigation, writing original draft, review and editing. S.L: investigation, writing review and editing, B.O: investigation, writing review and editing. G.S. writing review and editing. M.T.H: Conceptualization, supervision, funding acquisition, writing original draft, review and editing.

## Data availability

Code and data generated are available on github https://github.com/michhulin/Hulinlab/. RNA sequencing reads are available on NCBI Sequence Read Archive under BioProject PRJNA1419900.

## Supplementary materials

### Supplementary tables

Table S1: Plasmids used in this study.

Table S2: List of PCR primers used in this study.

Table S3: Results of VipTree and vConTACT3 analysis of prophages across PSSC

Table S4: Functional gene predictions in the PamPP1 prophage

Table S5: RNA sequencing statistics

Table S6: Normalized read counts and differentially expressed gene statistics

Table S7: Normalized read counts for genes of interest

Table S8: T3E prediction in the Pam IR 634 genome

### Supplementary Figure legends

Figure S1: The PSSC contains diverse prophages and the PamPP1 prophage is conserved across the complex. A: VipTree phylogeny of PSSC prophages with other *Pseudomonas* phages belonging to the Caudoviricetes. PSSC prophages are highlighted with blue circles and *Pseudomonas* phages with purple, The PamPP1 prophage from our work is highlighted in orange and with an arrow. The first circle on the heatmap shows groups identified using hierarchical clustering of the VipTree phylogeny, whilst the second circle shows the corresponding vConTACT3 families. The third circle shows which PSSC species (based on 95% average nucleotide identity) each prophage was identified within. The fourth circle shows whether each prophage was predicted to be incomplete, intact or questionable using PHASTEST. B: A core genome phylogenetic tree of 1669 strains from across the PSSC. Tips are highlighted by *P. syringae* species at the 95% average nucleotide identity level (colors the same as in A). The heatmap shows the presence of VipTree prophage groups, group 26 is highlighted with an orange arrow

Figure S2: Genes within the PamPP1 prophage region appear conserved within other pathovars in the *P. amygdali* clade, indicating that the prophage was gained in a common ancestor. (A) Alignment of the pv. *morsprunorum* conserved PamPP1 prophage region, anchored around *hopAR1*. (B) Alignment of pv. *eriobotryae* against the Pam IR 634 reference *hopAR1* prophage region anchored around *hopAR1*. (C) Alignment of pv. *myricae* against Pam IR 634 reference *hopAR1* prophage region. Genes are colored by blue for bacterial roles, green for prophage and *hopAR1* and pseudogenized *hopBK1* effectors in red, hypothetical proteins in black and unaligned proteins in grey highlighting the conservation of genes within the prophage region

Figure S3: Bacterial population growth assays comparing IR 634:ΔP, IR 634:Δ*hopAR1* and the IR 634 wildtype in leaf tissue. (A) Populations at 0 days post inoculation (B) Populations at 7 days post inoculation. For all graphs, significant differences tested via ANOVA (P<0.05 Tukey post-hoc testing) are shown by the compact letter display groupings above.

Figure S4: Removal of the PamPP1 prophage region does not cause altered growth in a range of media. (A) Growth measured every 15 mins over 24 h in 200 µL in KB with orbital shaking. (B) Growth measured every 15 mins over 24 h in 200 µL in LB with orbital shaking. (C) Growth measured at 0, 6, 9 24, 48 and 72 h in 100 mL of HIM with orbital shaking at 180 RPM. Graphs show standard deviation error bars (n=3) at each time point. No significant statistical differences were seen between the wildtype and mutant (calculated via ANOVA of the Area Under the Curve values).

Figure S5: RNA-seq samples show clear clustering based on strain and time with deviation growing between IR 634 and IR 634:ΔP over the course of 25 h induction in HIM. (A) Principal component analysis of all 24 RNA-seq samples (2 strains, 4 time points, 3 replicates per time point) using the variance stabilizing transformation of the DESeq2 count data. (B) Distanced based heatmap of all 24 RNA-seq samples using the variance stabilizing transformation of the DESeq2 count data.

Figure S6: Heatmaps of z-scores showing differentially expressed genes with an adjusted P < 0.05 at each time point. Samples are clustered by cladogram similarity of the DEG content showing that IR 634 and IR 634:ΔP consistently cluster with their replicates over time. Genes are also clustered by cladograms and likewise show clear clustering of downregulated genes vs upregulated genes between the three replicates at each time point. (A) Heatmap of DEGs in KB media prior to induction. (B) Heatmaps of DEGs at 0 h induction in HIM. (C) Heatmaps of DEGs at 5 h induction in HIM. (D) Heatmaps of DEGs at 25 h induction in HIM.

Figure S7: RNA-seq analysis shows no major differentially expressed genes outside of the PamPP1 prophage region between IR 634 and IR 634:ΔP when grown in KB. The analysis is performed for samples grown to an OD_600_ 0.2 in KB prior to induction. There were 41 genes with an adjp < 0.05, of which 5 are found outside of the prophage region. The analysis shows the MA plot (left) expressing the differential expression against the expression level of all genes for IR 634:ΔP against IR 634, genes with a padj < 0.05 are shown in blue. The EnchancedVolcano plot analysis (middle) shows genes with a log2FC >2 highlighted in green, thos with a p-value <10e-6 are shown in blue and genes displaying both those values are shown in red. Non-significant genes are shown in grey. Significantly differentially regulated genes of interest are labelled based on their locus tag. The Local Network Clusters (right) identified through string analysis against the proteome of IR 634 (STRG0A01MPX) terms are grouped with a similarity of ≥ 0.8, sorted by gene count and merged with a similarity ≥ 0.6 for genes with a padj of < 0.05.

Figure S8: Upset plot showing the shared differentially expressed genes between samples grouped by time and up or downregulated. The left graph in red shows the total number of genes associated to samples based on up or down regulated. Linkages between samples are shown to the right and the number of genes identified in that linkage shown in the graph above. Interesting connections include the 33 downregulated genes shared between all samples that account for the prophage region and 19 genes shared between the upregulated genes at 0 h, 5 h, and 25 h in HIM of which 17 are related to the T3SS.

Figure S9: Analysis of genes associated with important pathways identified through phenotypic analysis shows differentially regulated genes of interest. (A) gene counts of upstream T3SS regulators based on the literature identified through blast searching in Geneious Prime. Those labelled as “like” were additional distant homologs. (B) Gene counts of chemotaxis associated genes from genome annotations. (C) gene counts of type IV associated proteins from genome annotations. (D) gene counts of flagellar associated proteins from genome annotations. Plots were made using the normalized read counts from DEseq2 analysis, data is displayed as a normalized count scale by row of 0 to 1, then clustered based on expression patterns.

Figure S10: Standard curves for IR 634 and IR 634:ΔP HA tagged proteins in western blotting at 24 h in HIM shows higher production of T3SS associated proteins in IR 634:ΔP compared to the wildtype. For each western blot a serial dilution of the HA tagged protein of interest was performed from undiluted to 1:8 dilution, the chemiluminescent plot was then imported into ImageJ and the pixel density of the region measured, and the background pixel density used as a blank to produce an adjusted signal which was plotted against the concentration presented as a percentage (undiluted = 100%, 1:8 = 6.25%).

