## Supplementary figures and images for "It takes two: A Widespread Temperate Bacteriophage Contributes to Regulation of the Type III Secretion System in *Pseudomonas syringae*"

### Figure S1

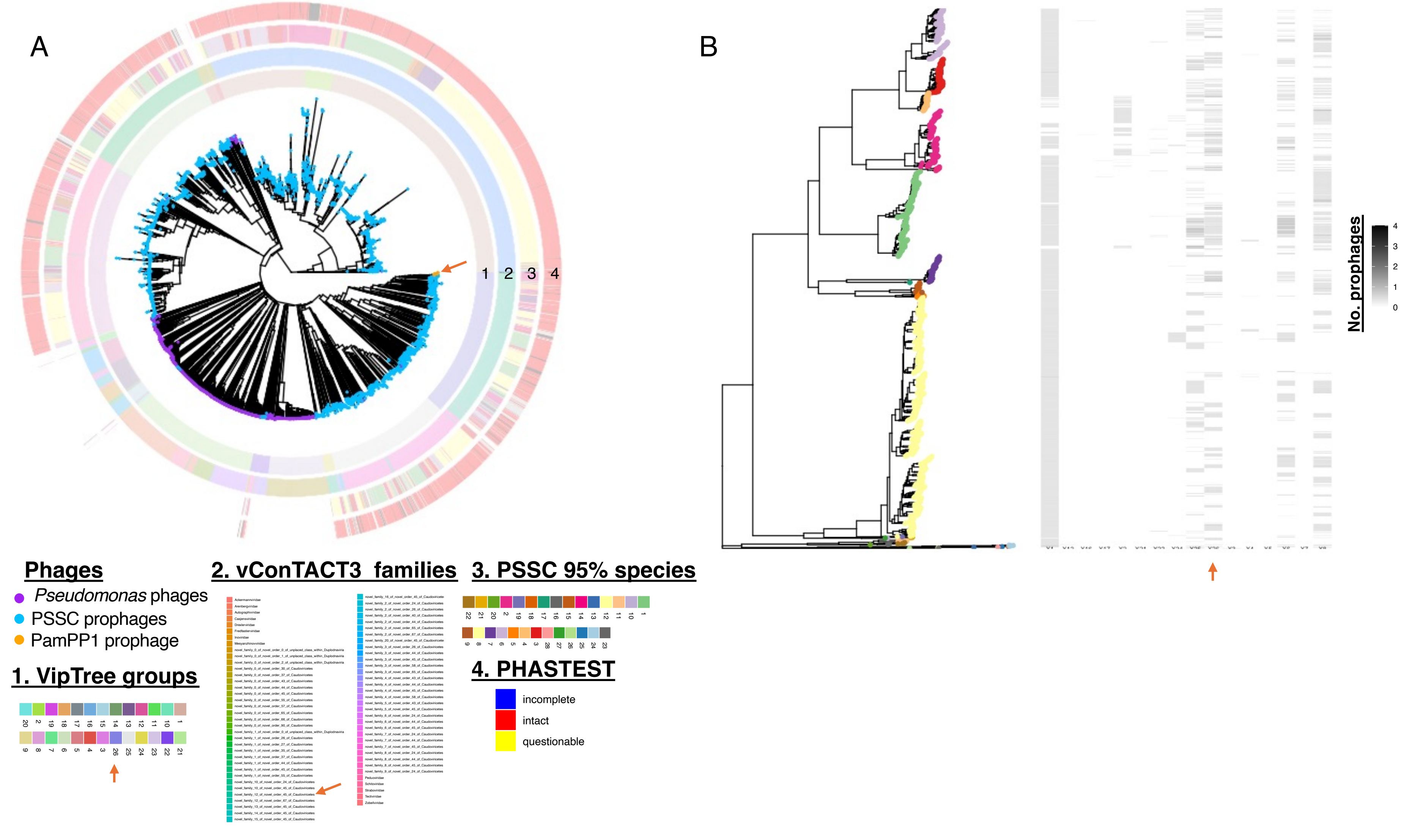

### Figure S2

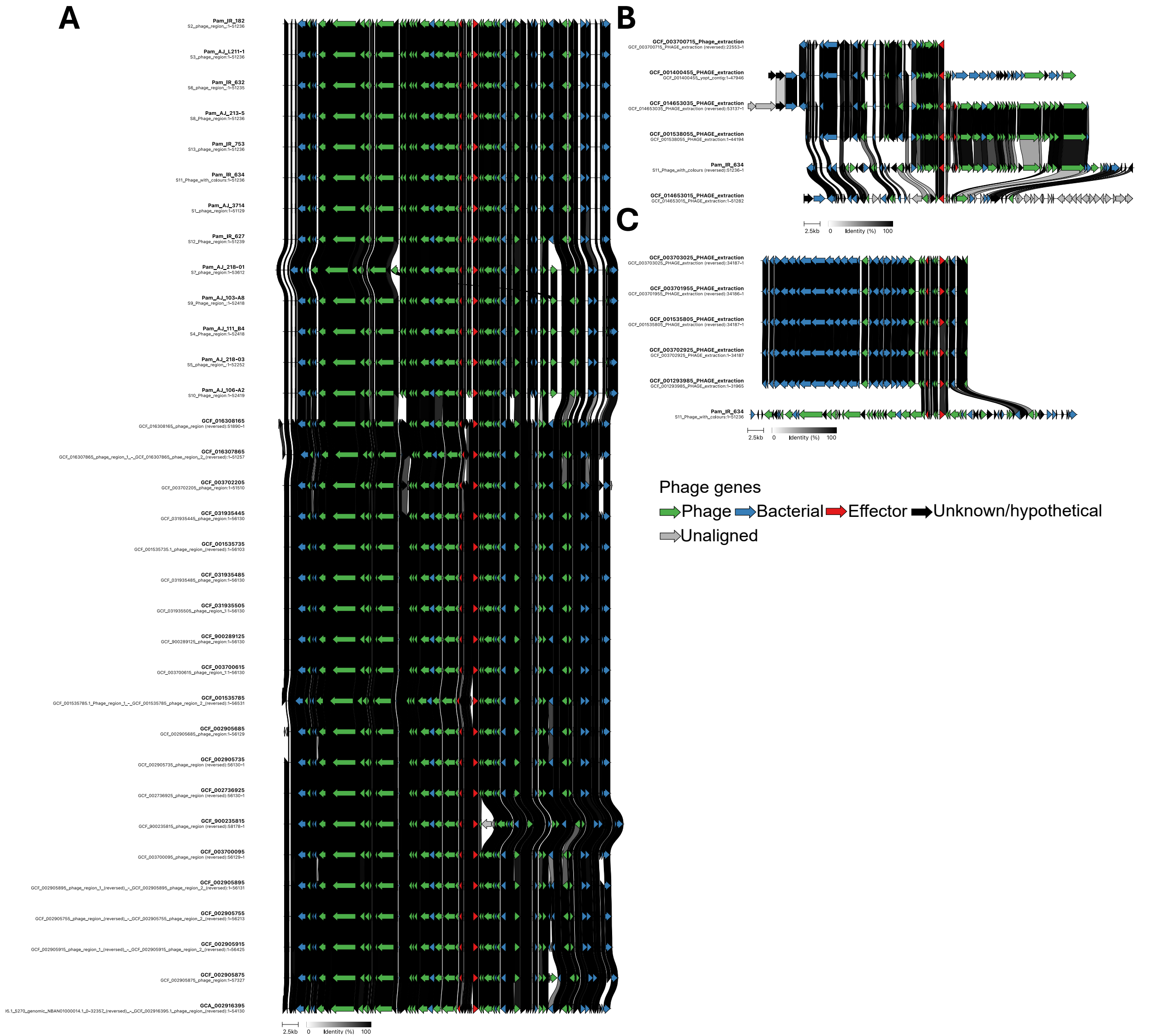

### Figure S3

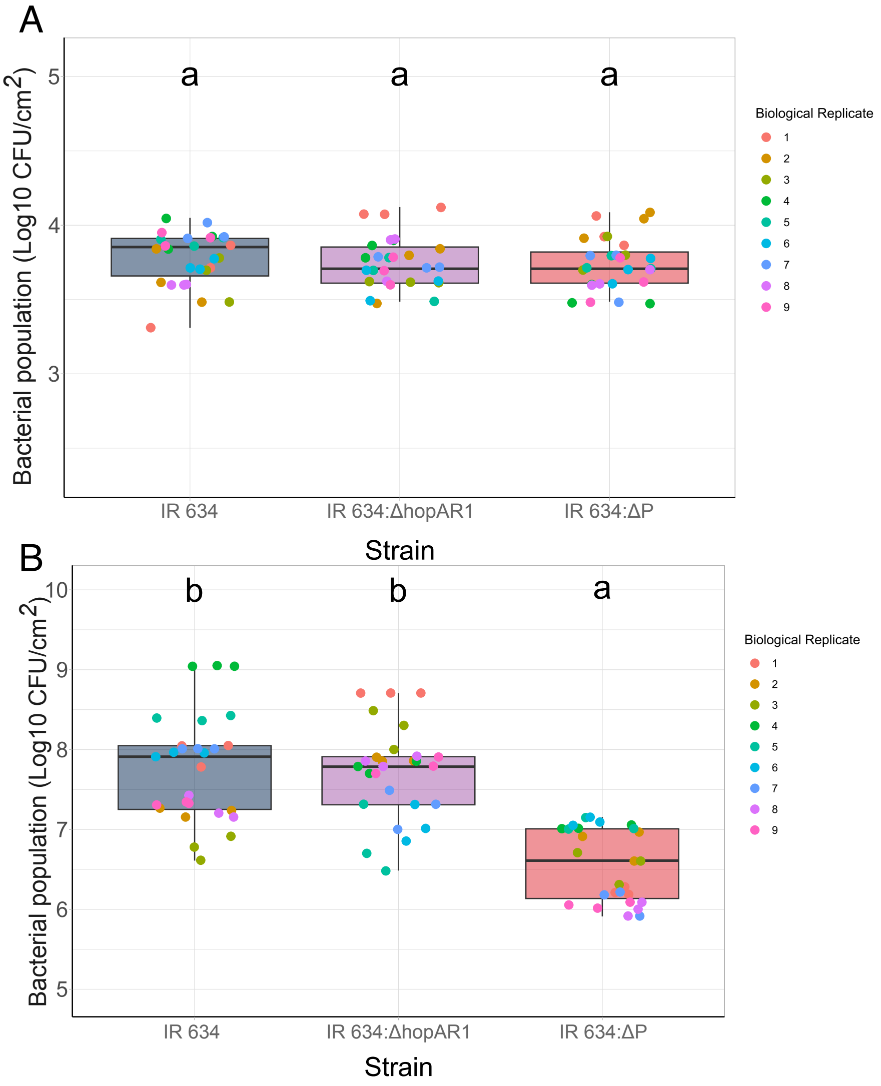

### Figure S4

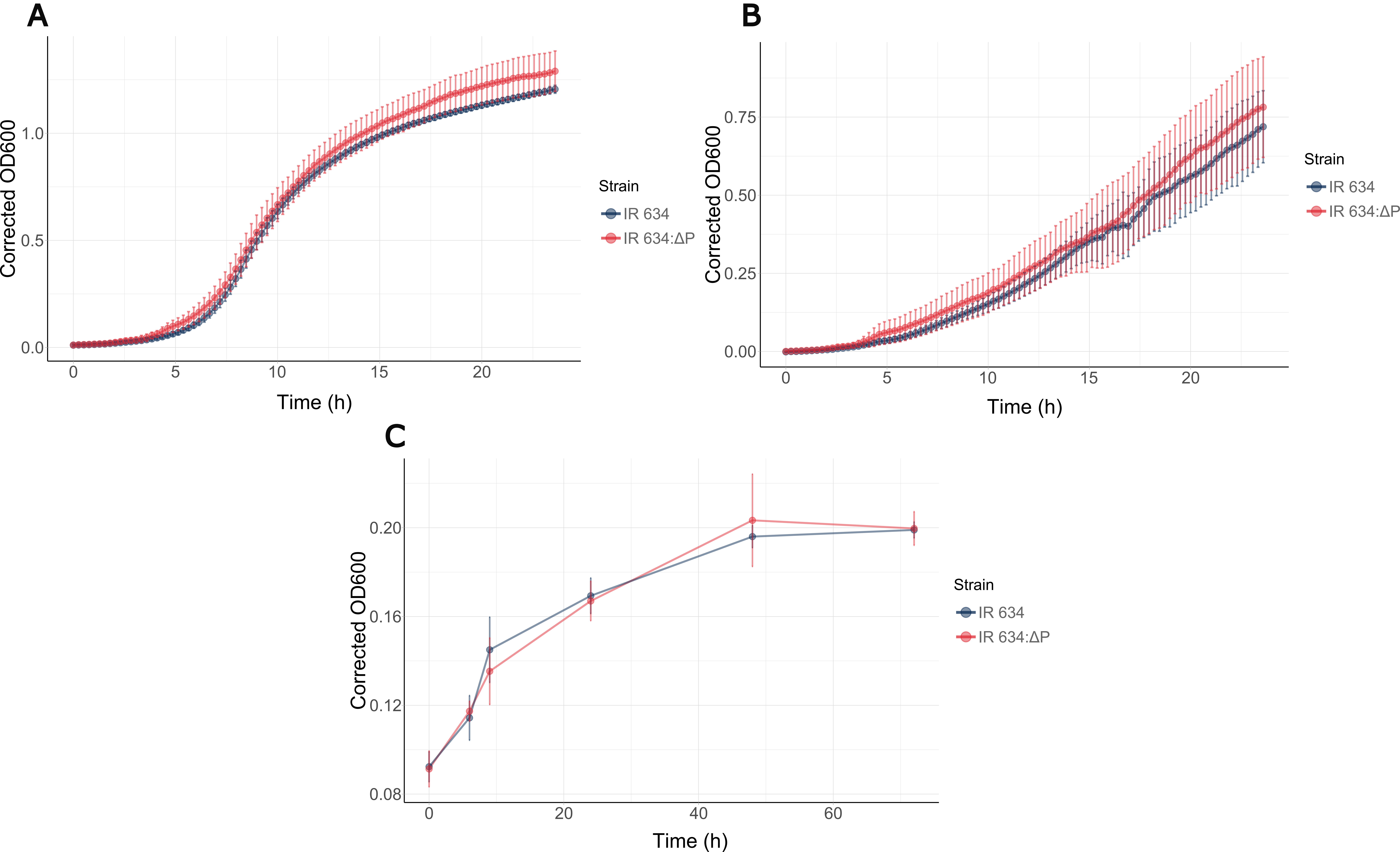

### Figure S5

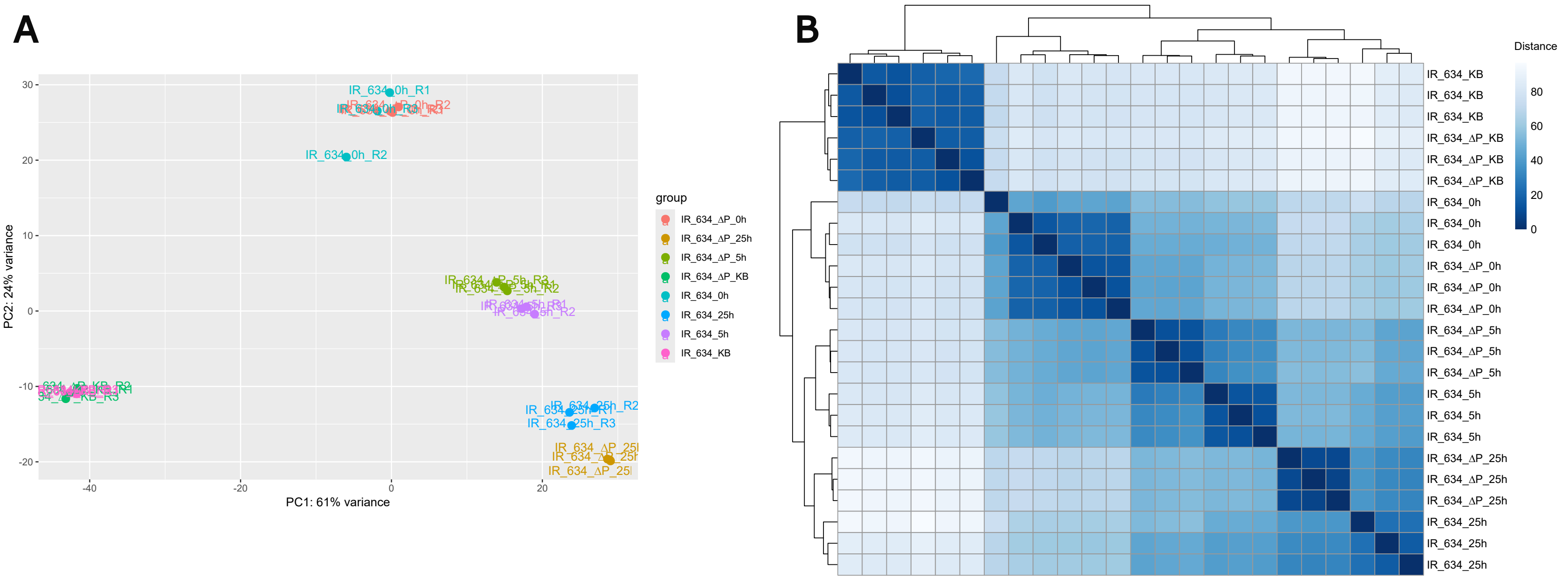

### Figure S6

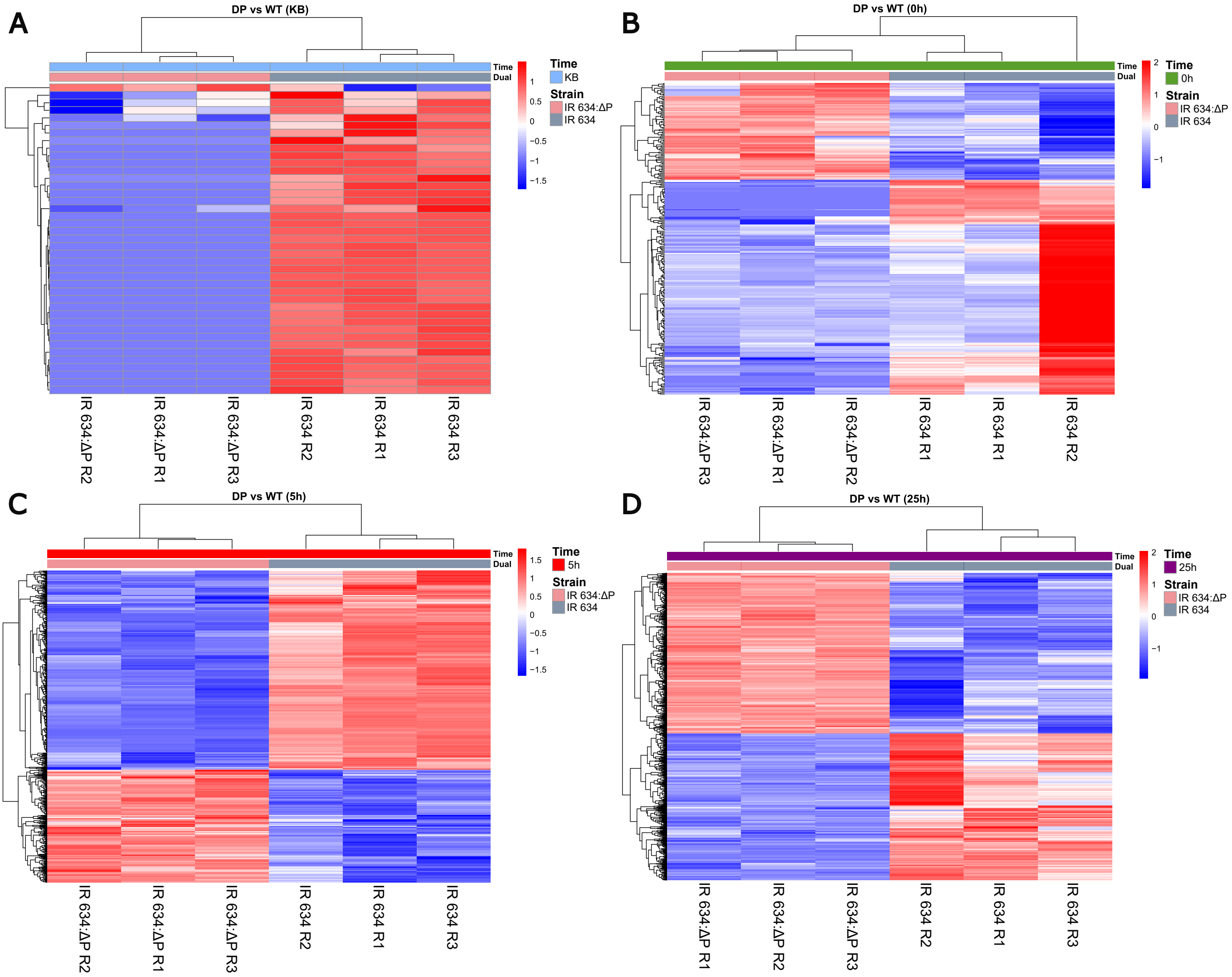

### Figure S7

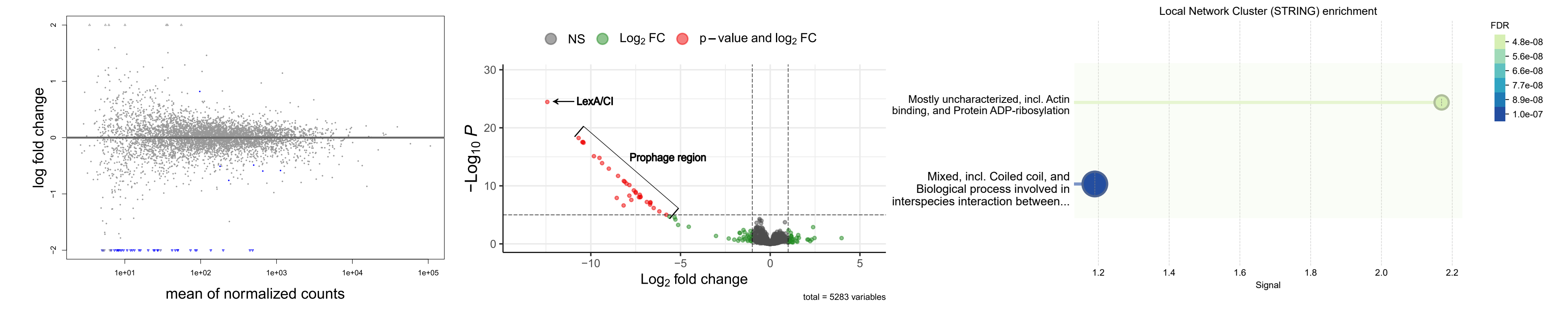

### Figure S8

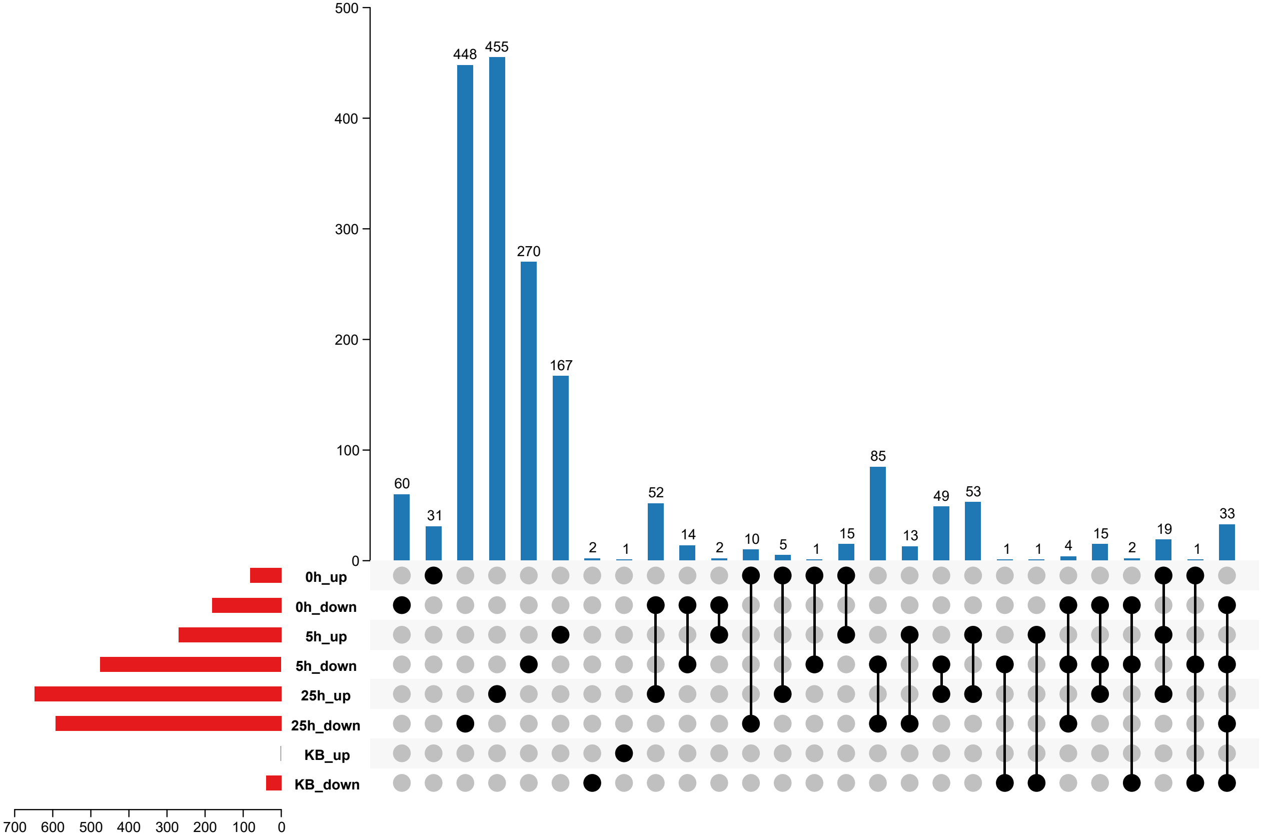

### Figure S9

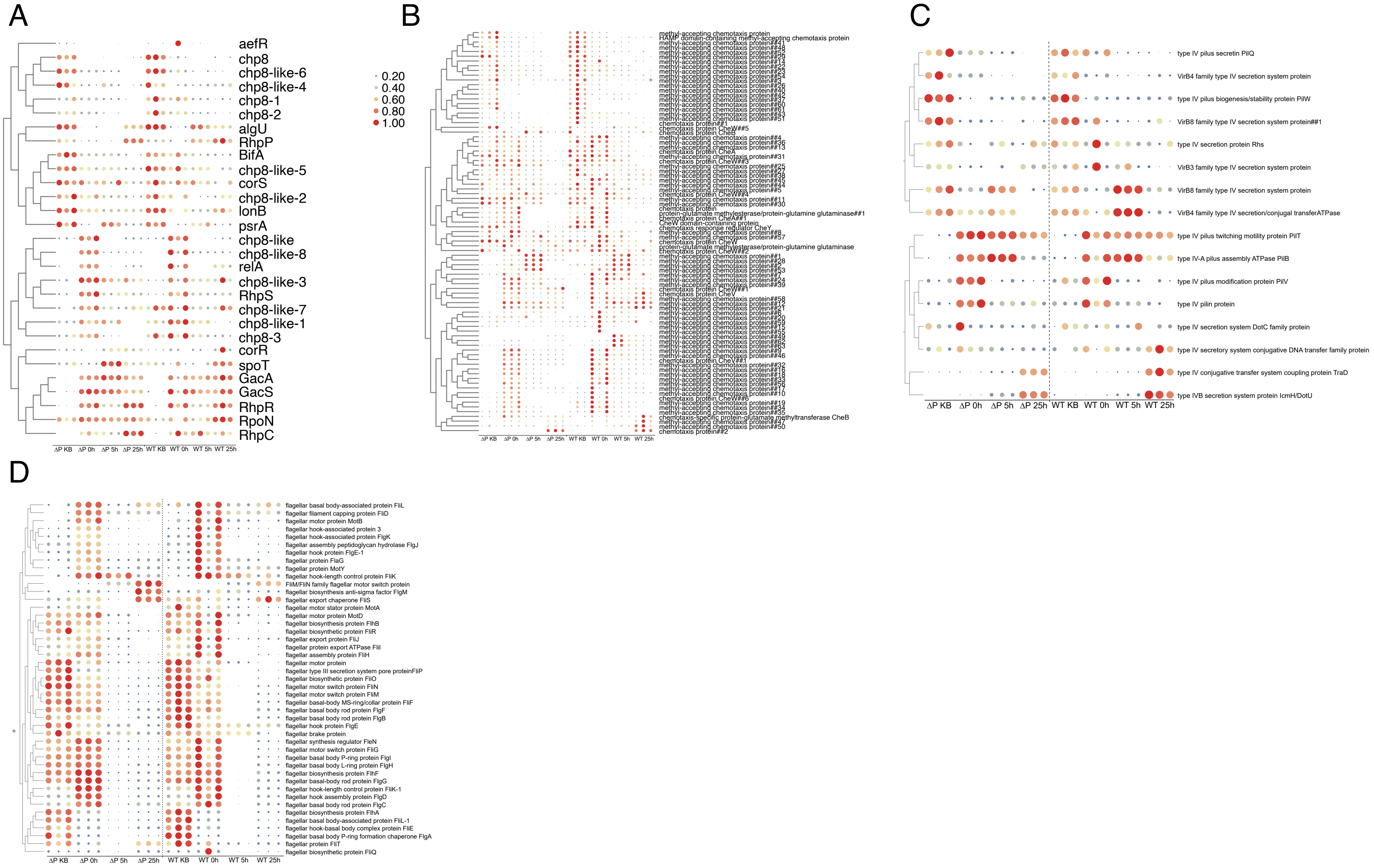

### Figure S10

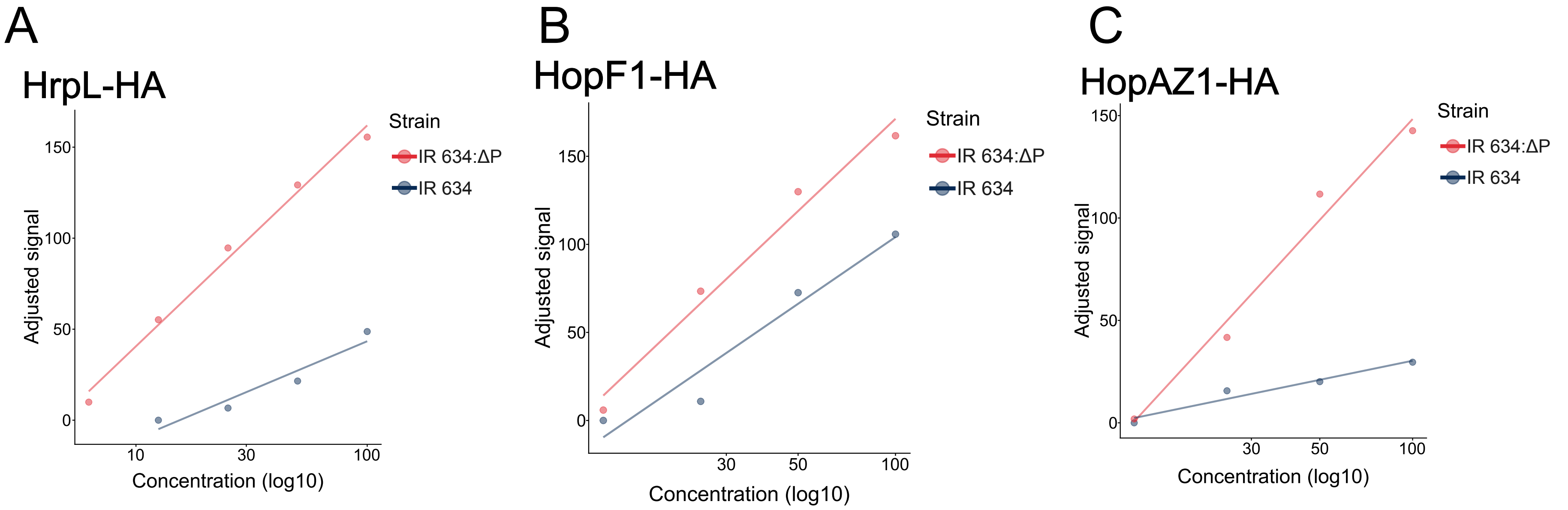
