## Supplementary material for "It takes two: A Widespread Temperate Bacteriophage Contributes to Regulation of the Type III Secretion System in *Pseudomonas syringae*": Table S1

| **Plasmid** | **Description** | **Features** | **Source** |
| --- | --- | --- | --- |
| pK18msB | Broad host range bacterial gene deletion/replacement vector | Suc^S^, Km^R^ | [60] |
| pRK2013 | Triparental mating helper plasmid | Km^R^,Tra+, Mob+, Col El replicon | [90] |
| pUCP20TK | Expression vector | Km^R^ | [62] |
| pK18msB-∆P-IR634 | For the deletion of the prophage region within *Pseudomonas amygdali* pv. *morsprunorum* | Suc^S^, Km^R^ | This study |
| pK18msB-∆*hopAR1*- IR634 | For the deletion of T3SS effector *hopAR1* within *Pseudomonas amygdali* pv. *morsprunorum* | Suc^S^, Km^R^ | This study |
| pK18msB-∆*hrpL*-IR634 | For the deletion of T3SS regulator RNA polymerase subunit sigma (*hrpL*) within *Pseudomonas amygdali* pv. *morsprunorum* | Suc^S^, Km^R^ | This study |
| pUCP20 -*hopAZ1*-HA | Hemagglutinin tagged T3E *hopAZ1* expressed under its native promoter | Km^R^ | This study |
| pUCP20 -*hopF*-HA | Hemagglutinin tagged T3E *hopF* expressed under its native promoter | Km^R^ | This study |
| pUCP20 -*hrpL*-HA | Hemagglutinin tagged T3SS gene RNA polymerase subunit sigma (*hrpL*) expressed under its native promoter | Km^R^ | This study |
| pUCP20 -*hrpA*-HA | Hemagglutinin tagged T3SS gene *hrpA* expressed under its native promoter | Km^R^ | This study |
