## Supplementary material for "It takes two: A Widespread Temperate Bacteriophage Contributes to Regulation of the Type III Secretion System in *Pseudomonas syringae*": Table S2

| **Primer ID** | **Sequence** | **Restriction site** | **Vector** | **DNA source** | **Primer purpose** |
| --- | --- | --- | --- | --- | --- |
| DM_0001_ppF1 | ACATGATTACGAATTATTCCCGGCGAAACAACC | EcoR1 | PK18msB | IR634 | Removal of PamPP1 prophage |
| DM_0002_ppR1 | AAAGCCCCGCAGATTAACGTGACCTCGGCCTCCACTATTC | None | PK18msB | IR634 |  |
| DM_0003_ppF2 | GAATAGTGGAGGCCGAGGTCACGTTAATCTGCGGGGCTTT | None | PK18msB | IR634 |  |
| DM_0004_ppR2 | CGACTCTAGAGGATCCGATCATGGCCTGGGTTG | BamH1 | PK18msB | IR634 |  |
| DM_0040_HopAR1del_F1 | ACATGATTACGAATTGCCAGGCCATCCACTCAATAAG | EcoR1 | PK18msB | IR634 | Deletion of *hopAR1* gene |
| DM_0041_HopAR1del_R1 | ATCGCTGGGGTCTTTCGTTTTACCTCGAATTTTTCCCGTTACG | None | PK18msB | IR634 |  |
| DM_0042_HopAR1del_F2 | CGTAACGGGAAAAATTCGAGGTAAAACGAAAGACCCCAGCGAT | None | PK18msB | IR634 |  |
| DM_0043_HopAR1del_R2 | CGACTCTAGAGGATCCCATGATGACTCGCAACACGC | BamH1 | PK18msB | IR634 | Deletion of *hrpL* gene |
| DM_137_hrpL_up_F | CTATGACATGATTACGAATTTTGGCGCTCATCAACAACTG | EcoR1 | PK18msB | IR634 |  |
| DM_138_hrpL_up_R | AGATATCCACGGGCTTACCTTAATTTAATGGTGTG | None | PK18msB | IR634 |  |
| DM_139_hrpL_down_F | AGGTAAGCCCGTGGATATCTGTCTGGAACCAACTC | None | PK18msB | IR634 |  |
| DM_140_hrpL_down_R | CAGGTCGACTCTAGAGGATCGGGCGACCATCGGATCCC | BamH1 | PK18msB | IR634 |  |
| DM_141_HA_hopAZ1_F | ACATGATTACGAATTTTAAGCGTAATCTGGAACATCG | EcoR1 | pUCP20TK | Synthetic fragment | Addition of HA tag to *hopAZ1* effector |
| DM_142_HA_hopAZ1_R | CGACTCTAGAGGATCCGAAACTGATCAGACAGCATGG | BamH1 | pUCP20TK | Synthetic fragment |  |
| DM_141_HA_hopF1_F | ACATGATTACGAATTTTAAGCGTAATCTGGAACATCG | EcoR1 | pUCP20TK | Synthetic fragment | Addition of HA tag to *hopF1* effector |
| DM_142_HA_hopF1_R | CGACTCTAGAGGATCATGCCGCTATGGAACTAAACC | BamH1 | pUCP20TK | Synthetic fragment |  |
| DM_145_HA_hrpL_F | ACATGATTACGAATTCTCAAGCGTAATCTGGAACATCG | EcoR1 | pUCP20TK | Synthetic fragment | Addition of HA tag to *hrpL* regulator |
| DM_146_HA_hrpL_R | CGACTCTAGAGGATCAGGACGGTTCTGAGCCTGG | BamH1 | pUCP20TK | Synthetic fragment |  |
